# Cellular and molecular basis of leptin resistance

**DOI:** 10.1101/2023.08.24.554526

**Authors:** Bowen Tan, Kristina Hedbacker, Leah Kelly, Zhaoyue Zhang, Ji-Dung Luo, Joshua D. Rabinowitz, Jeffrey M. Friedman

## Abstract

Obese humans and diet induced obese mice (DIO) have high leptin levels and fail to respond to the exogenous hormone suggesting that obesity is caused by leptin resistance, the pathogenesis of which is unknown. We found that leptin treatment reduced plasma levels of mTOR ligands leading us to hypothesize that mTOR activation might inhibit leptin signaling. Rapamycin, an mTOR inhibitor, reduced fat mass and increased leptin sensitivity in DIO mice but not in mice with defects in leptin signaling. Rapamycin restored leptin’s actions on POMC neurons but failed to reduce the weight of mice with defects in melanocortin signaling. mTOR activation in POMC neurons caused leptin resistance while POMC specific mutations in mTOR activators decreased the weight gain of DIO mice. Thus increased mTOR activity in POMC neurons is necessary and sufficient for the development of leptin resistance in DIO mice establishing a key pathogenic mechanism leading to obesity.

## INTRODUCTION

Obesity is the cardinal feature of the metabolic syndrome and a worldwide public health problem ^1^. Obesity develops when food intake exceeds energy expenditure leading to deposition of excess adipose tissue. In lean animals, adipose tissue mass is tightly controlled by the hormone leptin which functions as the afferent signal in a negative feedback loop that maintains energy balance. Leptin acts on key neural populations in hypothalamus to control food intake and mutations that alter signaling by leptin or melanocortins, downstream mediators of leptin action, cause extreme obesity in humans and rodents. Leptin reduces appetite by activating αMSH (POMC) expressing neurons in the arcuate nucleus of the hypothalamus and most of the known mutations that cause obesity alter the production of αMSH or its target, the MC4R G protein coupled receptor. Animals with leptin receptor or melanocortin mutations become hyperleptinemic and show a reduced response to exogenous leptin which are the hallmarks of a hormone resistance syndrome ^2,3^. These leptin resistant animals also show abrogated phosphorylation of the STAT3 transcription factor, a canonical biochemical effect of leptin signaling ^4–6^.

Leptin resistance can also be acquired when mice are fed a high-fat diet (HFD) leading to Diet Induced Obesity (DIO) but, in contrast to animals with mutations in the leptin-melanocortin axis, the pathogenesis of diet-induced leptin resistance has not been determined. Similar to DIO mice, most obese humans also develop leptin resistance and the response of DIO mice to anti-obesity therapies is highly predictive of a human response ^7,8^. Thus delineating a cause of leptin resistance in DIO mice would advance our understanding of the pathogenesis of obesity. Reversing leptin resistance could also have clinical implications particularly since leptin spares lean body mass in contrast to the new peptide-based therapeutics that can cause significant loss of lean mass ^9^.

Here we present a comprehensive set of physiologic, genetic and neurobiological studies showing that increased activity of the mTOR kinase in POMC neurons is necessary and sufficient for the development of leptin resistance and obesity in DIO mice.

## RESULTS

### A Metabolomic Screen for Biomarkers for Leptin Resistance

Animals and humans with low leptin levels lose weight on leptin therapy and we initially sought to identify acute biomarkers of a leptin response as a possible means for identifying responders even before weight loss develops ^3,7,8^. Toward that end, we collected plasma from leptin-sensitive and leptin-resistant animals before and after leptin treatment among four groups: wild-type mice fed chow (WT-Chow) or a high-fat diet (WT-DIO), and *ob/ob* mice fed chow (OB-Chow) or a high-fat diet (OB-HFD). In this study we controlled for food intake by pair feeding the OB-Chow group to chow fed mice, WT-Chow, group and by pair feeding the OB-HFD group to DIO mice, WT-DIO group, all for 18 weeks (Fig. 1a). Growth curves revealed that the WT-Chow group remained lean while the other three groups became increasingly obese (Fig. 1b, c, S1e, f). Plasma was collected from all the animals after vehicle treatment at week 18, and again after leptin treatment at week 19 (Fig. 1a). As expected, acute leptin treatment decreased food intake and body weight in the WT-Chow, OB-Chow and OB-HFD groups but did not affect the leptin-resistant WT-DIO group (Fig. 1d). Plasma metabolites were measured and analyzed by first normalizing all four groups of samples at each timepoint, followed by hierarchical clustering based on levels of metabolites after leptin treatment, (see heatmaps, Fig. 1e, S1a, b). These clustered heatmaps revealed a specific cluster of metabolites that were higher in the DIO group after leptin treatment compared to the levels after leptin treatment of WT-Chow, OB-Chow and OB-HFD mice. (Fig. 1e, Fig. S1a, b). We next identified pairwise, same-trend alterations in leptin-sensitive and resistant WT-DIO mice by comparing metabolite levels before and after leptin treatment. As previously reported ^10^, leptin decreased plasma triglycerides in leptin-sensitive mice but not in WT-DIO mice (Fig. 1f). We also found an increased level of several phosphatidic acids and glucosyl ceramides after leptin treatment of WT-DIO mice but these metabolites were unchanged in the other three groups (Fig. 1h, i), raising the possibility that leptin prevented the accumulation of these metabolites in the sensitive animals. Finally, we found a significant decrease of plasma leucine and methionine levels in leptin-sensitive but not in leptin resistant WT-DIO mice (Fig. 1g, S1c). Leucine and methionine are canonical activators of the mTOR pathway, and phosphatidic acids and ceramides can also modulate the PI3K-mTOR pathway ^11–13^. Our finding that leptin sensitivity was inversely associated with plasma levels of these mTOR ligands led us to hypothesize that mTOR activation might diminish leptin sensitivity in DIO mice. We evaluated this possibility by treating DIO with rapamycin, a specific mTOR inhibitor.

**Figure 1.**
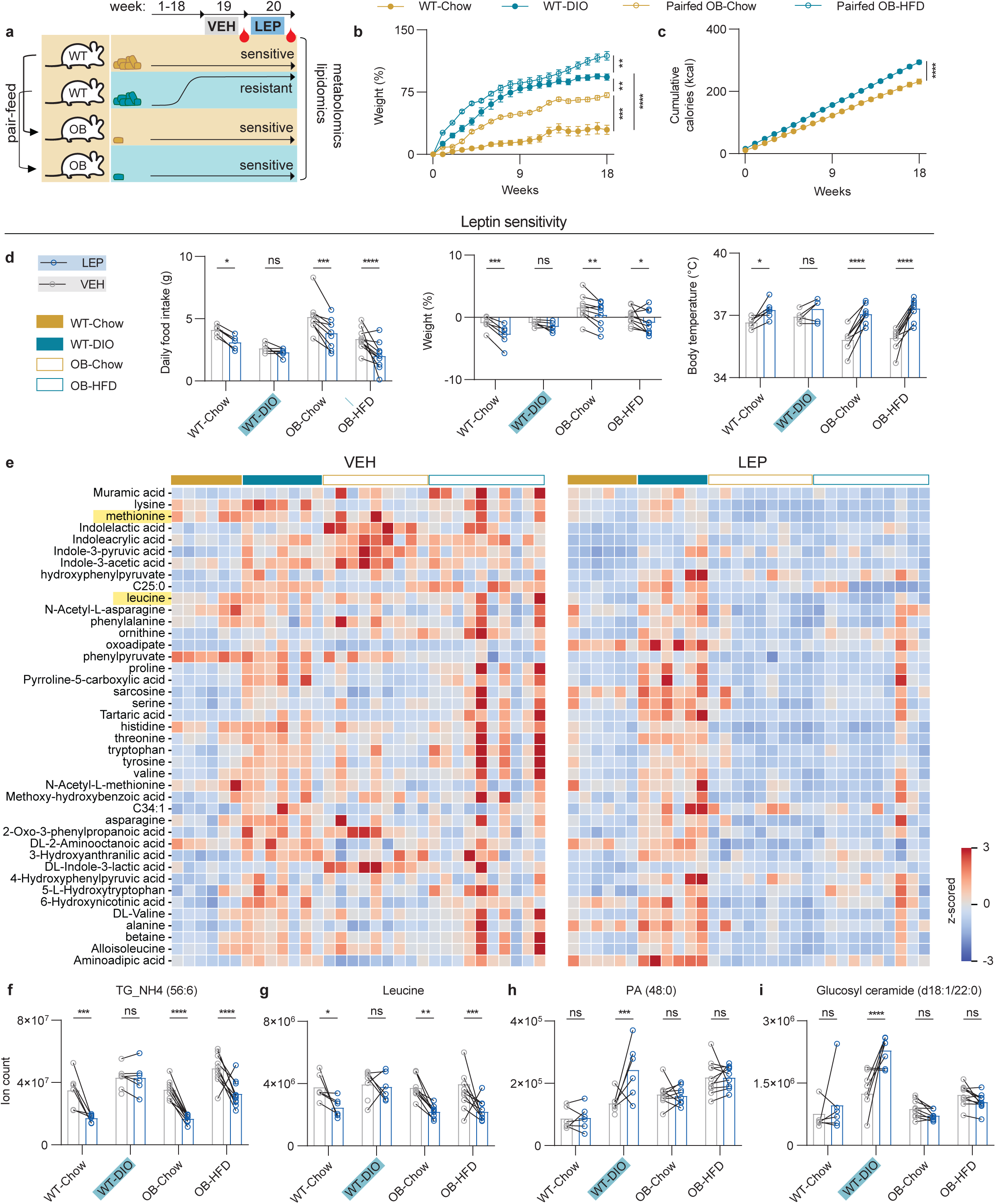
Metabolomic and lipidomic profiling of plasma metabolites associated with leptin resistance. (a) Schematic of *ob/ob* (OB) animals pairfed to WT animals for 18 weeks fed a chow or HFD. At 19^th^ week, mice were i.p. injected with vehicle (VEH) every 12 hours for 24 hours followed by blood collection. At 20^th^ week, mice were i.p. injected with 12.5 mg/kg leptin (LEP) every 12 hours for 24 hours followed by blood collection. (b) Time-course percentage of body weight relative to starting point (week 0). (c) Cumulative calories consumed (kcal) of WT mice fed a chow vs. HFD over 18 weeks. (Two-way ANOVA, with Tukey’s multiple comparisons, n=6, 6, 9, 10 for WT-Chow, WT-DIO, OB-Chow, OB-HFD, respectively). (d) Comparisons of 24-hour food intake, 24-hour % weight, and body temperature post VEH vs. post LEP (Two-way ANOVA, with Fisher’s LSD comparisons, n=6, 6, 9, 10 for WT-Chow, WT-DIO, OB-Chow, OB-HFD respectively). (e) Heatmap of the clustered metabolites levels higher in DIO post LEP vs. VEH. Comparisons of representative metabolites levels (ion count) post VEH vs. post LEP. (f) TG_NH4(56:6), (g) Leucine, (h) PA (48:0) and (i) Glucosyl ceramide (d18:1/22:0). (Two-way ANOVA, with Fisher’s LSD comparisons; n=6 for WT-Chow, n=6, 7 for DIO that received VEH vs. LEP, n=9 for OB-Chow, n=10 for OB-HFD). All error bars represent mean ± SEM. ns, not significant, *P < 0.05, **P < 0.01, ***P < 0.001, ****P < 0.0001. Additional information is included in Fig. S1.

### Rapamycin Reduces Obesity by Sensitizing DIO Mice to Leptin

DIO mice were treated with daily intraperitoneal (i.p.) injections of rapamycin (RAP) or vehicle (VEH) for 10 weeks. The rapamycin-treated DIO mice showed a decrease in daily and cumulative food intake (RAP: 149.1 ± 4.1 g vs. VEH: 186.6 ± 5.5 g; p<0.001), body weight (RAP: –27.6 ± 2.0 % vs. VEH: 3.7 ± 2.1 %; p<0.0001), a marked decrease of fat mass (RAP: –11.1 ± 1.4 g vs. VEH: 4.2 ± 1.2 g; p<0.0001), a small decrease in lean mass (RAP: –4.8 ± 0.4 vs. VEH: –1.0 ± 0.3; p=0.0514), and a highly significant decrease of fat-to-lean ratio (RAP: –0.3 ± 0.1 a.u. vs. VEH: 0.2 ± 0.0 a.u.; p<0.0001) (Fig. 2a-d, S2j). A similar decrease in lean mass after rapamycin has been previously reported, due to inhibition of protein synthesis ^14,15^. Thus, similar to the effects of leptin on leptin-sensitive animals ^2^, rapamycin reduced food intake, body weight and preferentially reduced fat mass relative lean mass in leptin-resistant DIO animals (Fig. 2c, d). This similarity between rapamycin treatment of DIO mice to leptin treatment of lean mice raised the possibility that rapamycin might sensitize animals to their endogenous leptin. We tested this by pre-treating DIO mice with rapamycin and then analyzing the response to injections of exogenous leptin.

**Figure 2.**
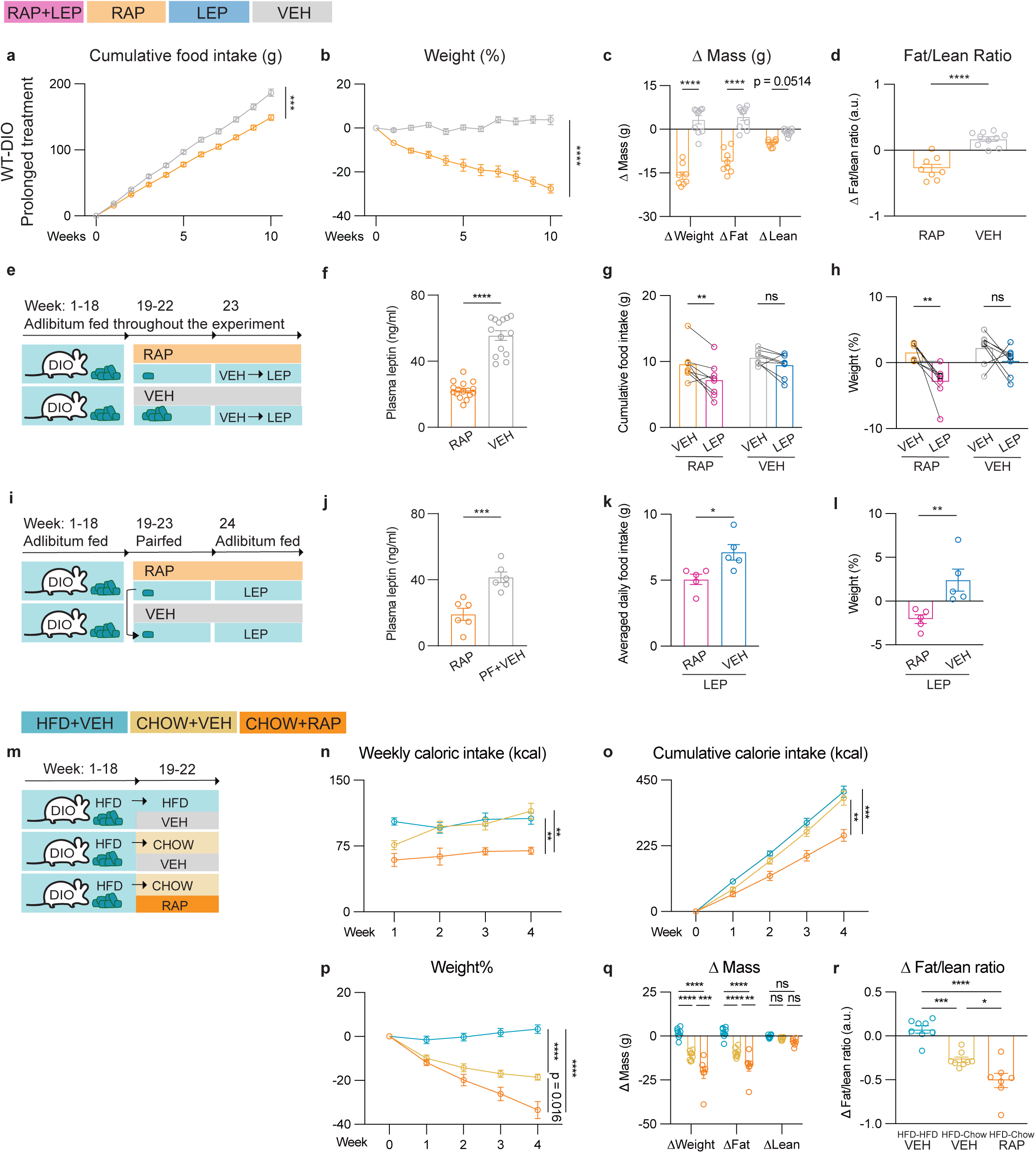
Rapamycin reduces food intake, fat mass, fat-to-lean ratio in DIO mice and increases their responses to the exogenous leptin. WT-DIO Prolonged treatment: DIO mice were treated with daily 2 mg/kg i.p. rapamycin (RAP) or vehicle (VEH) for 10 weeks. (a) Cumulative food intake, and (b) Weight, (c) Δ Mass of weight, fat and lean tissue, (d) Δ Fat/Lean Ratio (n = 8, 10 respectively; Two-way ANOVA, with Šidák’s multiple comparisons for Fig. 2a-c; Two-tailed Student’s t-tests for Fig. 2d). (e) Schematic of DIO mice that were treated with daily i.p. injections of 2 mg/kg Rapamycin (RAP) vs. vehicle (VEH) followed by a leptin sensitivity test. Each group was treated with i.p. injections of 2 mg/kg leptin (LEP) twice a day for 3 days followed by i.p. injections of vehicle (VEH) twice a day for 3 days. (f) Plasma leptin in DIO mice after 3-week RAP vs. VEH treatment prior to a leptin sensitivity test (n = 16, 14 for RAP, VEH, respectively; two-tailed Student’s t-tests). (g) Cumulative food intake and (h) Weight during leptin sensitivity test (n = 8, 8 for each group; two-way ANOVA, with Šidák’s multiple comparisons). (i) Schematic of DIO mice that were treated with daily i.p. injections of RAP vs. VEH. The group of mice treated with VEH were pairfed to the group of mice treated with RAP for 5 weeks. Leptin sensitivity test was conducted after 5-week treatment. For the leptin sensitivity test, each group of mice were treated with i.p. injections of 2 mg/kg LEP twice daily for 3 days followed by i.p. injections of VEH twice daily for 3 days. Both groups of mice had adlibitum access to food during the test. Additional information is included in Fig. S5a-d. (j) Plasma leptin in DIO mice after 3-week RAP vs. Pairfed-VEH treatment prior to the leptin sensitivity test (n = 6, 6 for each group; Two-tailed Student’s t-tests). (k) Cumulative food intake and (l) Weight of RAP vs. pairfed VEH-treated DIO mice were treated with LEP (n = 6, 6; Two-tailed Mann-Whitney tests). (m) Schematic of DIO mice that either remained on HFD with daily i.p. VEH-injections or transferred to chow-feed and given daily RAP vs. VEH injections. (n) Weekly caloric food intake (o) cumulative calorie food intake, (p) Weight, (q) Δ Mass of weight, fat and lean tissue, and (r) Δ Fat/Lean Ratio (n=8, 7, 8 respectively; Two-way ANOVA, with Tukey’s multiple comparisons for Fig. 2n-r; One-way ANOVA, with Tukey’s multiple comparisons for Fig. 2r). Additional information is included in Fig S5a,b.

We treated DIO mice with rapamycin for three weeks, after which they were treated with either leptin or vehicle for three days (Fig. 2e). Subsequent leptin injections elicited a further decrease in food intake (Fig. 2g, 9.6 ± 0.9 g vs. 7.2 ± 0.9 g; p<0.01), body weight (Fig. 2h, 1.6 ± 0.5 % vs. –2.9 ± 0.9 %; p<0.01), and a specific decrease in fat mass (Fig. S5a, b) in DIO mice pre-treated with rapamycin compared to those pre-treated with vehicle. Because pre-treatment with rapamycin reduced the weight of DIO mice, and hence leptin levels (Fig. 2f), a separate group of DIO mice were treated with vehicle for five weeks while being pair-fed to the ad libitum fed rapamycin-treated group on a HFD (Fig. 2i). We then tested leptin sensitivity in these pair-fed animals. The response to leptin in the pair-fed DIO mice was significantly reduced compared to the rapamycin-treated group, with higher food intake (7.1 ± 0.6 g vs. 5.1 ± 0.4 g; p<0.05) and weight (+2.4 ± 1.3 % vs. –2.1 ± 0.5 %; p<0.01) (Fig. 2k, l, S5c, d). Consistent with their lower weight, the plasma leptin levels were lower in rapamycin-treated DIO mice compared to the pair-fed vehicle group (19.1 ± 3.7 ng/ml vs. 41.5 ± ng/ml; p<0.01) (Fig. 2j). We also evaluated metabolism of the DIO animals using indirect calorimetry-enabled metabolic cages and found that rapamycin-treated DIO mice exhibited a decrease of respiratory exchange ratio (RER) (RAP 0.76 ± 0.00 RER vs. VEH 0.78 ± 0.01 RER; p<0.01), and an increase in energy expenditure (RAP 1.2 ± 0.1 kcal/g vs. VEH 1.0 ± 0.0 kcal/g; p<0.05) compared to the vehicle group (Fig. S5e-h).

As a further control, we fed DIO mice a high-fat diet for 18 weeks after which the diet was changed to chow while simultaneously administering either rapamycin or vehicle (Fig. 2m). Consistent with prior reports ^16,17^, when the vehicle-treated mice were switched from a HFD to chow, there was a significant reduction of body weight of ∼20% (from 61.4 ± 1.0 g to 48.1 ± 3.0 g; p=0.007; Fig. 2p) This was also associated with a decrease of food intake, fat mass and fat-to-lean ratio but this reduction plateaued by the fourth week at weights that were still considerably higher than chow fed controls that had never been fed a high fat diet (35.0 ± 1.3 g; Fig. S1e). In contrast DIO mice transferred to a chow diet while receiving rapamycin showed significantly greater weight loss, a greater reduction of fat mass, and a more sustained reduction in food intake than did the vehicle-treated group (from 58.2 ± 3.0 g to 40.0 ± 2.2 g; p< 0.0001, Fig. 2n-p). Overall, these data are consistent with the restoration of leptin’s effects after pre-treatment with rapamycin, and suggest that rapamycin, but not weight loss which develops after the mice are switched from a high fat diet to chow can re-sensitize DIO mice to their endogenous leptin. If true, rapamycin should have little or no effect on food intake or fat mass in animals with defects in leptin signaling. We tested this by comparing the effects of rapamycin on lean control mice and *ob/ob* and *db/db* mice.

Similar to 10 weeks of rapamycin treatment, DIO mice treated with rapamycin for 14 days showed a significantly decreased food intake (RAP: 32.4 ± 1.6 g vs. VEH: 39.7 ± 1.9 g; p < 0.001), body weight (RAP: –10.2 ± 1.1 % vs. VEH: 0.1 ± 1.1 %; p < 0.0001), fat mass (RAP: –5.2 ± 1.3 vs. VEH: 0.5 ± 0.6; p < 0.001), fat-to-lean ratio (RAP: –0.1 ± 0.1 a.u. vs. VEH: 0.1 ± 0.0 a.u, p < 0.05) with a small decrease in lean mass (Fig. 3a-d, S2i). In contrast, rapamycin did not decrease food intake or fat-to-lean ratio in *ob/ob* and *db/db* mice fed a chow or a high-fat diet (Fig. 3i-p, S2a-h). Similar to its effect on DIO mice, rapamycin caused a modest weight loss, that was limited to a loss of lean mass in *ob/ob* (Δweight: –3.9 ± 0.7 g, Δlean mass: –2.8 ± 0.2 g) and *db/db* (Δweight: – 6.0 ± 2.1 g, Δlean mass: –5.6 ± 0.7 g) mice fed a high-fat diet (Fig. 3i-p, S2a-h,l,m). Thus animals with defects in leptin signaling failed to show a decrease of food intake or fat mass with rapamycin treatment. A 14-day course of rapamycin also had little or no effect on food intake, body weight or fat mass in lean, chow-fed mice which are leptin sensitive with low endogenous hormone levels (Fig. 3e-h, S2k). We next tested whether leptin synergizes with rapamycin in these animals as well as in aged obese mice which also have relatively low leptin levels compared to DIO mice.

**Figure 3.**
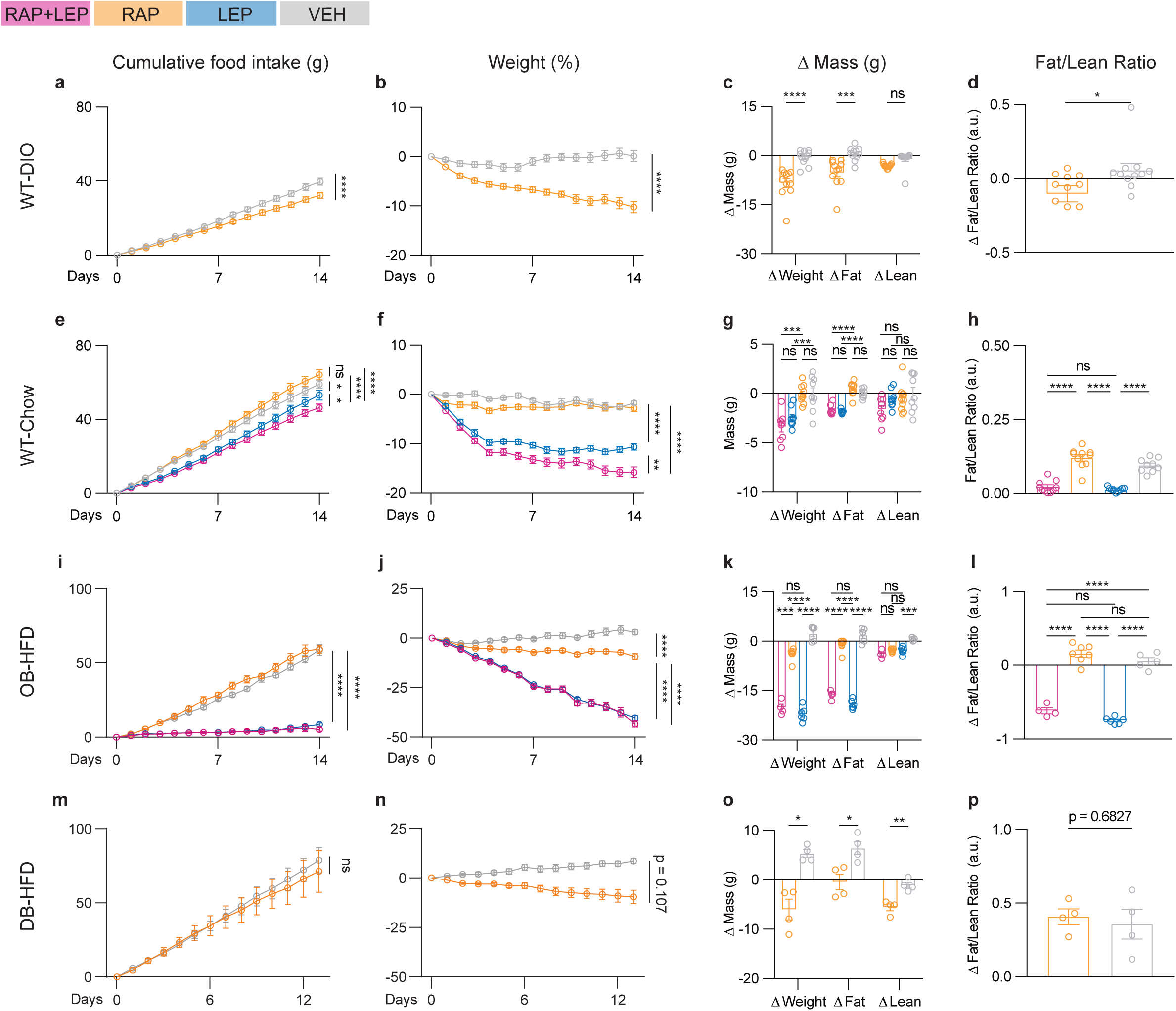
Rapamycin reduces food intake, fat mass, fat-to-lean ratio in DIO mice but not in *ob/ob* or *db/db* mice. WT-DIO: DIO mice were treated with RAP or VEH daily for 14 days. (a) Cumulative food intake, and (b) Weight, (c) Δ Mass of weight, fat and lean tissue and (d) Δ Fat/Lean Ratio. (n = 11, 11 for each group; two-way ANOVA, with Šidák’s multiple comparisons for Fig. 2a-c; Two-tailed Student’s t-tests for Fig. 2d). WT-Chow: Chow-fed lean mice were treated with daily RAP plus 600 ng/hr leptin (LEP), RAP plus VEH, LEP plus VEH, VEH plus VEH for 14 days. (e) Cumulative food intake, and (f) Weight, (g) Δ Mass of weight, fat and lean tissue and (h) Fat/lean Ratio at day 14. (n = 12, 14, 14, 13 for RAP+LEP, RAP, LEP, VEH, respectively; Two-way ANOVA with Tukey’s multiple comparisons for Fig. 2e-g; One-way ANOVA with Tukey’s multiple comparisons for Fig. 2h). OB-HFD: *ob/ob* mice fed a HFD were treated with 2 mg/kg daily RAP plus 300 ng/hr leptin (LEP), RAP plus VEH, VEH plus LEP, VEH plus VEH for 14 days. (i) Cumulative food intake, (j) Weight, (k) Δ Mass of weight, fat and lean tissue and (l) Fat/Lean Ratio (a.u.) at day 14. (n = 10, 13, 11, 10 for RAP+LEP, RAP, LEP, VEH, respectively; Two-way ANOVA with Tukey’s multiple comparisons for Fig. 2i-k; One-way ANOVA with Tukey’s multiple comparisons for Fig. 2l). DB-HFD: *db/db* mice fed a HFD were treated with daily i.p. injections of RAP (2 mg/kg) vs. VEH for 13 days. (m) Cumulative food intake, and (n) Weight, (o) Δ Mass of weight, fat and lean tissue and (p) Δ Fat/Lean Ratio (a.u.) measured at Day 0 and Day 13. (n = 4, 4, respectively; Two-way ANOVA, with Šidák’s multiple comparisons for Fig. 2m-o; Two-tailed Student’s t-tests for Fig. 2p). All error bars represent mean ± SEM. ns, not significant, *P < 0.05, **P < 0.01, ***P < 0.001, ****P < 0.0001. Additional information is included in Fig. S2, S3, S4.

We first compared the effects of high-dose (600 ng/hr) to low-dose leptin (150 ng/hr) with or without rapamycin in lean, chow-fed mice. As previously reported, the high dose of leptin resulted in a significantly greater decrease of weight and fat mass than did the lower leptin dose (Fig. S4a-d) ^3^. We next tested a combination of low-dose leptin (150 ng/hr) combined with rapamycin and found that the combination elicited a significantly greater reduction of food intake, body weight and fat mass than did low-dose leptin alone. Moreover, the final weight of the lean, chow-fed mice treated with low-dose leptin (150 ng/ml) plus rapamycin was comparable to that of mice treated with high-dose leptin (600 ng/hour) (low-dose LEP+RAP vs. high-dose LEP; Weight: –10.1 ± 1.0 % vs. –8.6 ± 2.4 %; p = ns, Food intake: 59.8 ± 5.9 g vs. 53.9 ± 3.4 g; p=ns, ΔFat mass: –1.2 ± 0.3 g vs. –1.0 ± 0.2 g; p = ns) (Fig. S4a-h) with both groups showing a similar decrease in adipose tissue mass relative to lean mass (Fig. S4c, g). These data show synergy between leptin and rapamycin in chow fed mice.

Aged mice also develop obesity, but have been reported to show only modestly increased leptin levels of ∼ 10 ng/ml compared to ∼ 5 ng/ml in lean mice ^18^. While aged mice weigh more than younger chow fed mice, 16-month-old animals were significantly less obese than DIO mice (Aged weight: 39.8 ± 0.9 g, n = 23; DIO weight: 53.9 ± 0.9 g, n = 19, p<0.0001. Aged fat: 24.4 ± 1.5 %, n = 14 vs. DIO fat: 45.1 ± 0.6 %; p<0.0001 (Fig. S4j, S4n, 3b, 3c). Consistent with their relatively low leptin levels, i.p. injections of rapamycin alone had no effect on food intake, body weight or fat mass compared to vehicle-treated mice. We next compared the effect of high-dose leptin (600 ng/ml) with or without rapamycin. Leptin alone resulted in a small but significant decrease of food intake, body weight, fat mass and fat-to-lean ratio (Fig. S4i-p). However, the addition of rapamycin to the leptin treatment significantly reduced food intake, body weight and fat mass (LEP+RAP: 46.9 ± 3.8 g vs. LEP: 57.3 ± 4.8 g; p<0.05), body weight (LEP+RAP: –13.8 ± 3.3 % vs. LEP: –5.4 ± 2.8 %; p<0.01), fat mass (LEP+RAP: –4.9 ± 0.5 g vs. LEP: –2.8 ± 0.5 g; p < 0.05) with no significant effect on lean mass (p = 0.32) (Fig. S4i-p).

Finally, we compared the effect of rapamycin with and without high-dose leptin in DIO mice. Consistent with their high baseline leptin levels (∼48 ng/ml, Fig. S4u), the food intake, body weight and fat mass of DIO mice was similar in animals treated with rapamycin plus leptin (600 ng/hr) vs. rapamycin alone (Fig. S4q-t). However, the addition of leptin to rapamycin did have a significant effect to improve glucose tolerance. Rapamycin has been shown to impair glucose tolerance while leptin has been shown to improve it ^19,20^. Consistent with previous studies, chronic rapamycin treatment increased plasma glucose levels in chow-fed lean mice, *ob/ob* and *db/db* mice (Fig. S3a,c,d) while a combination of leptin and rapamycin treatment lowered the baseline plasma glucose levels in WT-Chow, WT-DIO and OB-Chow animals (Fig. S3a,b,c). To further address the improvement of glucose tolerance by leptin, we performed glucose-tolerance tests in DIO and chow fed mice. In both cases, rapamycin alone worsened glucose tolerance, and the addition of exogenous leptin significantly improved it. This improvement was characterized by a lower peak glucose level in the chow fed mice, a reduction of the area under the curve in DIO and chow fed mice (WT-Chow: RAP 5.4 x 10^4^ ± 0.3 x 10^4^ a.u. vs. RAP+LEP 3.2 x 10^4^ ± 0.2 x 10^4^ a.u.; p<0.0001. DIO: RAP 9.3 x 10^4^ ± 0.9 x 10^4^ a.u. vs. RAP+LEP 6.9 x 10^4^ ± 0.5 x 10^4^ a.u.; p<0.05) and the normalization of plasma glucose after 90-120 minutes in both DIO and chow fed mice (Fig. S3 e-h). Thus, while chronic rapamycin treatment can impair glucose tolerance, the addition of exogenous leptin mitigates it.

In aggregate, these data suggest that rapamycin synergizes with exogenous leptin in chow fed and aged mice which have lower baseline hormone levels and can sensitize DIO animals to their endogenous leptin (Fig. S9a, b). We next set out to establish the mechanism responsible for the re-sensitization of leptin signaling in DIO mice after rapamycin treatment.

### The Cellular Target of Rapamycin-Mediated Leptin Sensitization

We began by mapping anatomic sites with increased mTOR activity in DIO mice by performing whole-brain imaging to map levels of phosphoS6 (pS6), an mTOR substrate and canonical marker for its activity, using the SHIELD brain clearing procedure ^21^. The hypothalamus showed the largest increase of pS6 levels in DIO mice compared to chow-fed mice (Fig. S6a, b). Numerous studies have shown that leptin regulates energy balance by modulating the activity of specific cell types in the hypothalamus, particularly the arcuate nucleus (ARC) ^6,22,23^. We thus assayed pS6 levels in the ARC of DIO mice before and after rapamycin treatment using IHC. Consistent with the SHIELD data, there were significantly elevated pS6 levels in the ARC of DIO mice compared to chow-fed lean mice and the increased levels in DIO mice were significantly reduced by rapamycin treatment (Fig. 4a, k, S6a, b).

**Figure 4.**
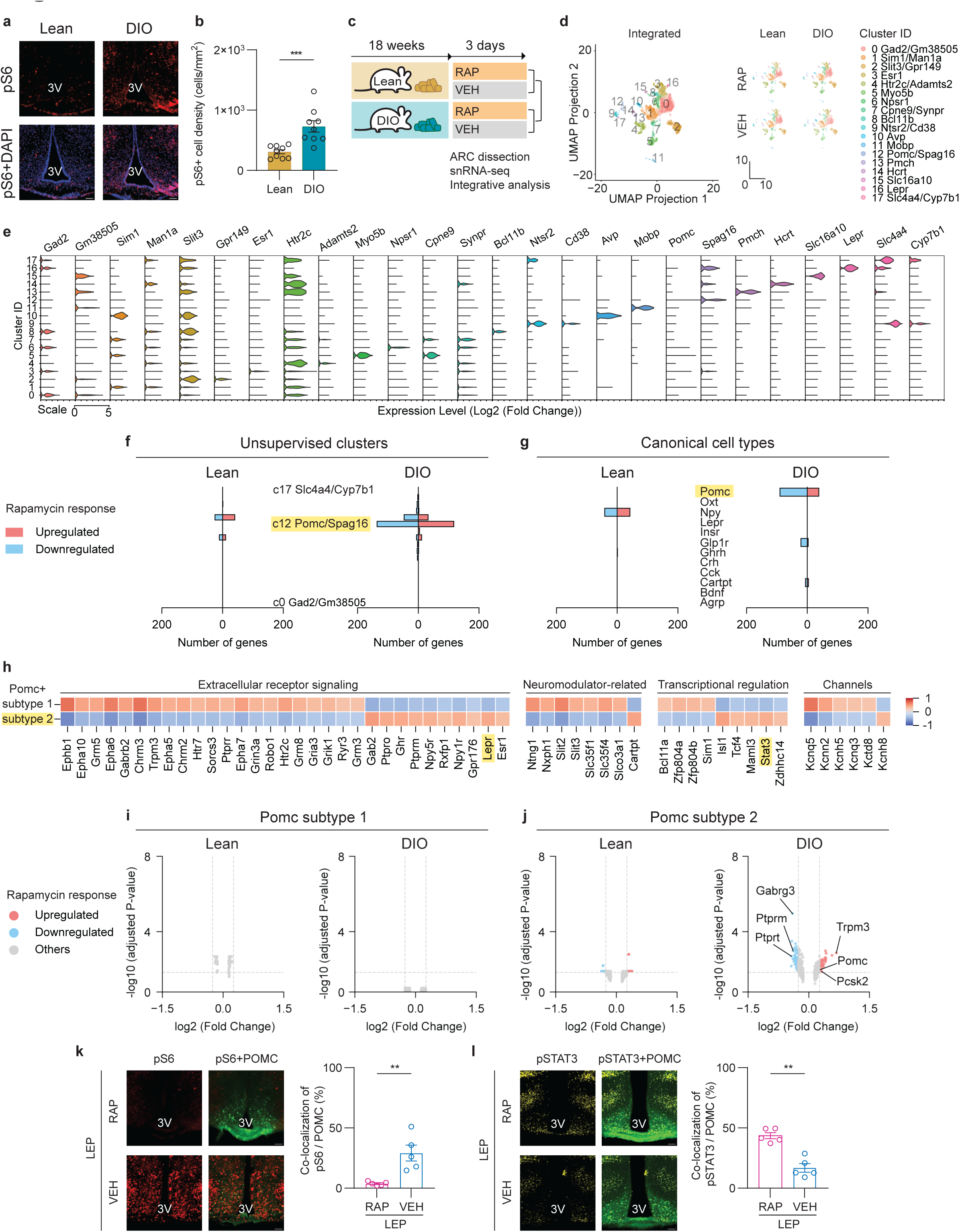
Single-nucleus RNA sequencing reveals that hypothalamic POMC neurons primarily respond to rapamycin in a leptin-signaling-dependent manner. (a) Immunohistochemical staining of pS6 levels, and DAPI, in DIO mice compared to chow-fed lean mice. (b) Quantification of pS6 densities in DIO mice compared to chow-fed lean mice (n = 9 sections from 3 mice for each group, two-tailed Mann-Whitney tests). (c) Schematic of chow-fed lean mice or DIO mice followed by 3-day daily i.p. injections of 2 mg/kg RAP vs. VEH. 4-hour after last treatment, ARC tissues were microdissected for snRNA-seq with integrated analysis across groups. (d) UMAP representations of 18 clusters of cell types identified and shared across all Lean-RAP, Lean-VEH, DIO-RAP, DIO-VEH groups (n = 4934, 5182, 5992, 4514 nuclei profiled from the ARC for each group). (e) Violin Plot depicting the expressions levels of enriched molecular markers for each cluster. (f, g) Quantification of total numbers of regulated genes by RAP vs. VEH across unsupervised clusters and canonical cell types. (h) A heatmap representing the enriched molecular markers for POMC subtypes that are shared across all groups. (i, j) Volcano plots showing differentially expressed genes in POMC subtypes in response to RAP vs. VEH between Lean and DIO groups. (k) Immunohistochemical staining of pS6 and POMC-GFP levels in DIOs that received 3-day RAP vs. VEH treatment, followed by acute leptin treatment (2 mg/kg, i.p. injections). Quantification of co-localization of pS6 and POMC-GFP (n = 5 sections from 3 mice for each group, two-tailed Mann-Whitney tests). (l) Immunohistochemical staining of pSTAT3 and POMC-GFP levels in DIOs that received 3-day RAP vs. VEH treatment, followed by acute leptin treatment (2 mg/kg, i.p. injections). Quantification of co-localization of pSTAT3 and POMC-GFP (n = 5 sections from 3 mice for each group, two-tailed Mann-Whitney tests). All error bars represent mean ± SEM. ns, not significant, *P < 0.05, **P < 0.01, ***P < 0.001, ****P < 0.0001. Additional information is included in Fig. S6.

We next performed single-nucleus RNA sequencing (snRNA-seq) of the ARC to molecularly profile the response of individual cells to 3 days of rapamycin treatment vs. vehicle in chow-fed lean, DIO and *ob/ob* mice (Fig. 4c, S6c). This short-term treatment did not significantly alter food intake or body weight (see Fig. 2). The ARC was microdissected and single-nuclei libraries were prepared and sequenced. We used UMAP to define 18 distinct cellular clusters shared among all of the groups (Fig. 4d) and generated a violin plot of enriched molecular markers for each of them (Fig. 4e). These analyses revealed that rapamycin treatment of DIO mice significantly altered gene expressions in only a single cluster: cluster 12, which was defined by two marker genes *Pomc* and *Spag16* (Fig. 4f). There were only minimal, non-significant changes in the expression of the genes in this cluster in rapamycin-treated chow fed mice and a set of opposite changes was observed in *ob/ob* mice (Fig. S6c-g). POMC neurons are a canonical target of leptin action and these data thus suggested that the subtype of POMC neurons in this cluster might be a cellular target of rapamycin in DIO mice. To further characterize the POMC clusters and other known populations, we analyzed gene expression in ARC neurons expressing canonical marker genes for other cell types controlling feeding behavior and metabolism including *Pomc+, Oxt+, Npy+, Lepr+, Insr+, Glp1r+, Ghrh+, Crh+, Cck+, Cartpt+, Bdnf+* and *Agrp+* neurons. Consistent with the analysis of genes expressed in cluster 12, *Pomc+* neurons from DIO mice showed the largest transcriptomic alterations in response to rapamycin. Prior reports have indicated there are at least two distinct subsets of POMC neurons expressing either *Lepr* or *Glp1r* ^24–28^. To further evaluate which population showed significant changes after rapamycin treatment, we sub-clustered the POMC neurons into two distinct populations: POMC subtype 2 included *Lepr*, *Stat3* and *Spag16,* while POMC subtype 1 included *Htr2c* (Fig. 4h). We found that only POMC subtype 2 showed significant transcriptional changes after rapamycin treatment of DIO mice, while POMC subtype 1 did not (Fig. 4h-j). Further analysis of differentially expressed genes in POMC subtype 2 neurons from DIO mice revealed that rapamycin decreased the expression of *Ptprm* and *Ptprt*, which encode enzymes that dephosphorylate pSTAT3 ^29^, a key component of the leptin signal transduction pathway and *Gabrg3*, a GABA receptor subunit previously shown to inhibit POMC neurons ^30,31^. Rapamycin also increased the expression of *Pomc*, the precursor of α-MSH, *Pcsk2*, a POMC processing enzyme, and *Trpm3*, a cation channel (Fig. 4j). The expression levels of these genes in POMC cluster subtype 2 were unchanged in neurons from rapamycin-treated chow fed lean mice (Fig. 4j) and rapamycin treatment of *ob/ob* mice suppressed the expression of *Pomc* and *Pcsk2* in *Pomc+* cell types (Fig. S6f, g). These findings raised the possibility that rapamycin restored leptin action by reducing the expression of genes that diminish leptin signal transduction and increasing the expression of genes that increase the levels of α-MSH, a key neuropeptide produced by POMC neurons. To assess this, we examined the effect of rapamycin on the levels of pSTAT3 and pS6 in POMC neurons using a POMC-eGFP transgenic mouse line ^6^. At baseline, pS6 levels were increased in POMC neurons of DIO mice and pSTAT3 was expressed at very low levels (Fig. 4k, l). Three-days of rapamycin treatment decreased pS6 levels in POMC neurons and significantly increased tpSTAT3 levels compared to the vehicle group (Fig. 4k, l). These data suggest that rapamycin reduced the weight of DIO mice by re-sensitizing POMC neurons to the high endogenous leptin levels in these animals.

We evaluated this further by pre-treating animals with rapamycin for 3 days and then analyzing the effect of leptin (100nM) on the activity of POMC neurons in hypothalamic slices. We first performed cell-attached baseline recordings from GFP-expressing neurons in brain slices prepared from untreated, chow-fed, lean POMC-eGFP (lean) and untreated DIO POMC-eGFP (the DIO mice were fed a HFD for a minimum of 16 weeks). We also compared the baseline activity of POMC neurons from DIO mice pretreated for 3 days with either vehicle (DIO+VEH) or rapamycin (DIO+RAP) injections (Fig. 5a). POMC neurons from DIO and DIO+VEH mice exhibited significantly reduced firing rates compared to neurons from lean and DIO+RAP mice. The average firing rates from each group are as follows: lean: 2.00 ± 0.42 Hz (n = 51), DIO: 0.91 ± 0.11 Hz (n = 154; p=0.007), DIO+VEH: 0.81 ± 0.20 Hz (n = 36; p=0.003), and DIO+RAP: 2.75 ± 0.64 Hz (n = 28) (Fig. 5b). The baseline firing rate of DIO+RAP neurons was significantly increased compared to DIO+VEH (p=0.0004) and untreated DIO mice (p<0.0001, Fig. 5b). We also found that there was a significantly smaller proportion of spontaneously spiking POMC neurons from DIO mice compared to lean mice (57.80% of the total for DIO, n = 154 vs. 90.2% for lean mice, n = 51, p<0.0001, Fig. 5c), as has previously been reported ^32^. In addition, DIO+RAP significantly increased the number of spiking neurons compared to DIO+VEH (47.22% for DIO-VEH, n = 36; vs. 82.14% for DIO-RAP, n = 28, p<0.0001) (Fig. 5c).

**Figure 5.**
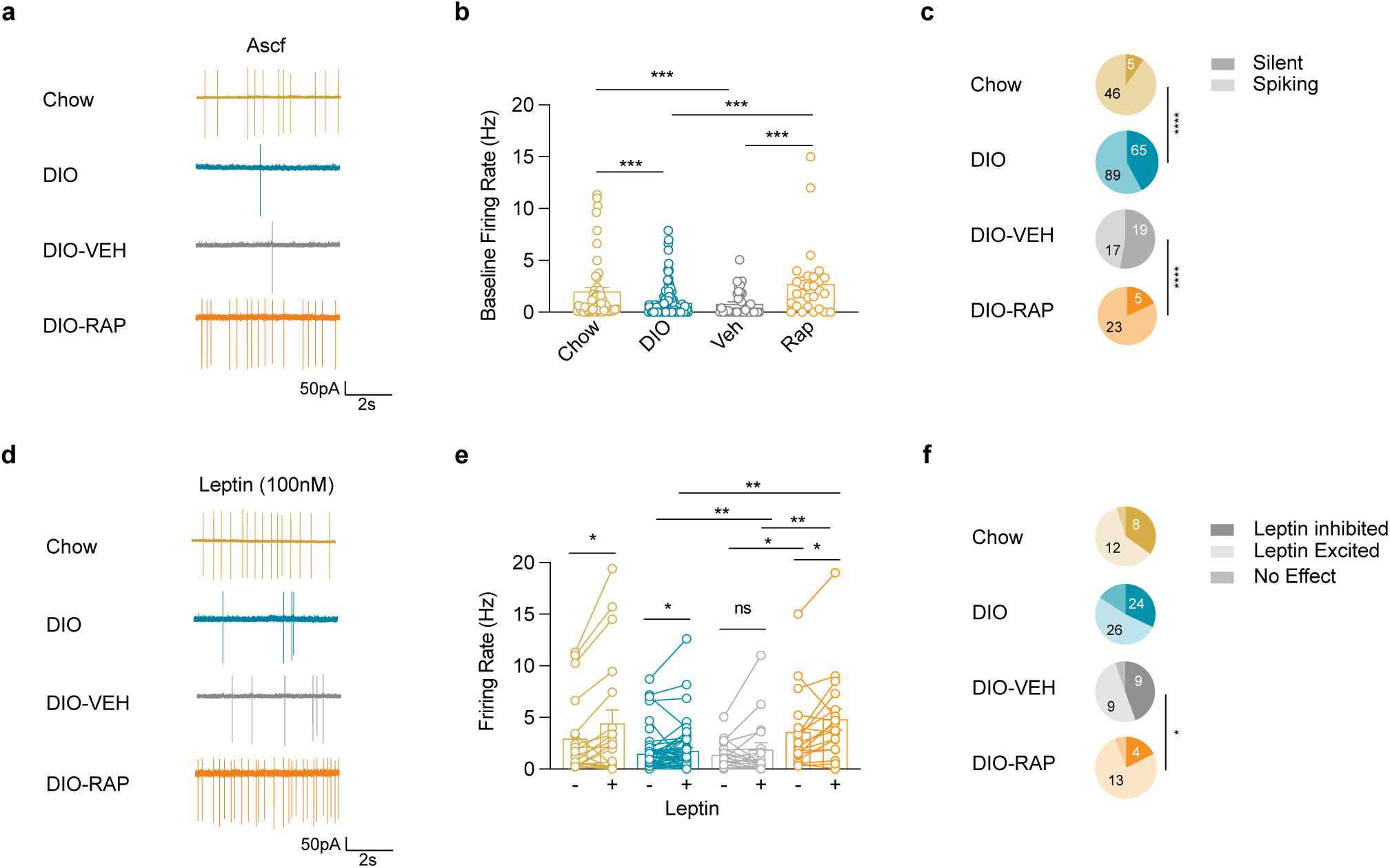
Rapamycin increases baseline firing rate and proportion of leptin-excited POMC neurons in DIO mice. (a) Cell-attached recordings from a chow-fed, DIO, DIO treated with vehicle (DIO-VEH) and from a DIO mouse treated with rapamycin (DIO-RAP) (top to bottom). Scale bar represents 50 pA for chow and DIO, 100 pA for vehicle and 20 pA for rapamycin and 2 seconds for all traces. (b) Baseline firing rate (Hz) of POMC neurons from recordsed from a chow-fed, DIO, DIO-VEH and DIO-RAP (left to right) (orange box represents i.p. injection). Firing rate is decreased in DIO (0.91 ± 0.11 Hz, n = 154; p = 0.007) and DIO-VEH (0.81 ± 0.20 Hz, n = 36; p = 0.003) compared to chow fed (2.00 ± 0.42 Hz, n = 51). Rapamycin rescues decreased firing rate in DIO (2.75 ± 0.64 Hz, n = 28) both compared to DIO with no treatment (p<0.0001) and DIO-VEH (p = 0.0004, all Mann-Whitney tests). (c) Percentage of spiking POMC neurons is decreased in DIO (57.80%, n = 154) and DIO-VEH (47.22%, n = 36) compared to chow (90.20% n = 51; both p < 0.0001, two-tailed binomial test). Rapamycin increases proportion of spiking POMC neurons (82.14%, n = 28) compared to DIO mice (p = 0.012) and DIO-VEH (p < 0.0001, both binomial test). (d) Cell-attached recordings from POMC-eGFP neurons in bath-applied leptin (100 nM, 20 mins) from chow-fed, DIO, DIO-VEH and DIO-RAP mice (top to bottom). Scale bar represents 50 pA for chow and DIO, 100pA for vehicle and 20 pA for rapamycin and 2 seconds for all traces. (e) POMC firing rates (Hz) in ASCF and bath-applied leptin (100 nM, 20 mins) recorded from chow-fed, DIO, DIO-VEH and DIO-RAP (left to right, orange box represents i.p. injection). Leptin increases firing rate in POMC neurons in chow-fed mice from 2.95 ± 0.34 to 4.41 ± 1.31 Hz (n = 20; p = 0.028), in DIO from 1.50 ± 0.29 to 1.77 ± 0.33 Hz (n = 50; p = 0.0238) and in DIO-RAP from 3.58 ± 0.88 to 4.81 ± 1.03 Hz (n = 17, p = 0.0208). Mean firing rate of POMC neurons of DIO-VEH mice does not change in the presence of leptin 1.38 ± 0.31 vs 1.87 ± 0.67 Hz (n = 18; p = 0.6095, all Wilcoxon matched pairs). Mean firing rate in leptin is higher in POMC neurons of DIO-RAP mice 4.81 ± 1.03 Hz (n = 17) compared to DIO-VEH mice 1.87 ± 0.67 Hz (n = 18; p=0.0047) and untreated DIO 1.77 ± 0.33 Hz (n = 50; p = 0.0002, both Mann-Whitney test). (f) Proportion of leptin-excited POMC neurons in chow-fed mice 60% (n = 20), DIO mice 52% (n = 50), DIO-VEH 50% (n = 18) and DIO-RAP mice 76.5% (n = 17) (from top to bottom). Rapamycin increases the proportion of leptin-excited neurons in DIO 76.5% (n = 17) compared to DIO-VEH (p = 0.0148, two-tailed binomial test).

We next evaluated the response of POMC neurons to leptin application *ex vivo* in slices prepared from the following groups; untreated lean animals, untreated DIO animals, mice pre-treated with vehicle for 3 days, (DIO+VEH), and mice pre-treated with rapamycin for 3 days, (DIO+RAP). Mean firing rates before and after leptin were calculated in 10 second bins (Fig. S9). Representative cell-attached traces are shown after leptin application (100nM, 20 mins) to POMC neurons in slices prepared from the four groups (Fig. 5d). Overall, leptin led to a significant 49 % increase of firing rate of POMC neurons from lean mice (2.95 ± 0.34 Hz to 4.41 ± 1.31 Hz; n = 20; p=0.028) (Fig. 5c), with a smaller 18 % increase of firing rate in neurons from DIO mice after application of leptin (1.50 ± 0.29 Hz to 1.77 ± 0.33 Hz; n = 50; p=0.0238). POMC neurons in slices prepared after rapamycin pre-treatment showed a 38% increase in firing rate after application of leptin (3.58 ± 0.88 Hz to 4.81 ± 1.03 Hz; n = 17, p=0.0208) (Fig. 5c) while the firing rate of DIO+VEH mice did not change significantly after application of leptin (1.38 ± 0.31 Hz vs 1.87 ± 0.67 Hz; n=18; p=0.6095) (Fig 5c). Rapamycin treatment also significantly increased the proportion of leptin-excited neurons to 76.5 % compared to 50 % (n = 18) of POMC neurons from vehicle-treated mice (n = 17; p=0.0148) (Fig. 5e).

### Melanocortin Signaling is Required for Rapamycin-Mediated Leptin Sensitization

Mice with POMC ablation have been previously shown to become obese ^33^, and if POMC neurons are the primary target of rapamycin, similar to *ob/ob* and *db/db* mice, these obese animals should not show a reduced fat mass after rapamycin treatment. We generated four groups of mice by stereotaxically delivering AAV5-hsyn-FLEX-mCherry or AAV5-hsyn-FLEX-dTA into the ARC of POMC-Cre mice (referred to as POMC-dTA vs. POMC-mCherry mice). Eight weeks post viral expression, the POMC-dTA mice were obese (Fig. S8a. POMC-dTA: 45.5 ± 2.0 g vs. POMC-mCherry: 29.0 ± 1.5 g) with higher fat mass (42.5 ± 1.3 % vs. 14.9 ± 2.3 %; p<0.0001) than the POMC-mCherry group which remained lean (Fig. S8a). Treatment of POMC-dTA animals with a high dose of leptin (600 ng/hr) had no effect on food intake, body weight or glucose tolerance confirming that these animals were leptin resistant (Fig. S8d-f). We then administered rapamycin vs. vehicle to POMC-dTA and POMC-mCherry mice (Fig. 6a) for 14 days. Rapamycin treatment of POMC-dTA mice did not significantly alter food intake (Fig. 6b. RAP: 66.5 ± 4.2 g vs. VEH: 78.2 ± 3.3 g; p=0.27), Δfat mass (Fig. 6d. –1.7 ± 0.3 g vs. 0.2 ± 0.7 g; n = 11, 13, p=0.12), or fat-to-lean ratio (Fig. 6e). Similar to the effects in *ob/ob* and *db/db* mice there was a small decrease in weight (RAP in dTA: –7.3 ± 1.1 % vs. VEH in dTA: 3.1 ± 2.1 %, p<0.01) secondary to a reduced lean mass compared to vehicle (RAP in dTA: –1.9 ± 0.3 g vs. VEH in dTA: 0.8 ± 0.5 g; p<0.01; Fig.6c, d, S6a). A combination of rapamycin and leptin also failed to alter food intake, body weight or glucose tolerance in the POMC-dTA animals (see Fig. S8g-i and a summary table in Fig. S9c).

**Figure 6.**
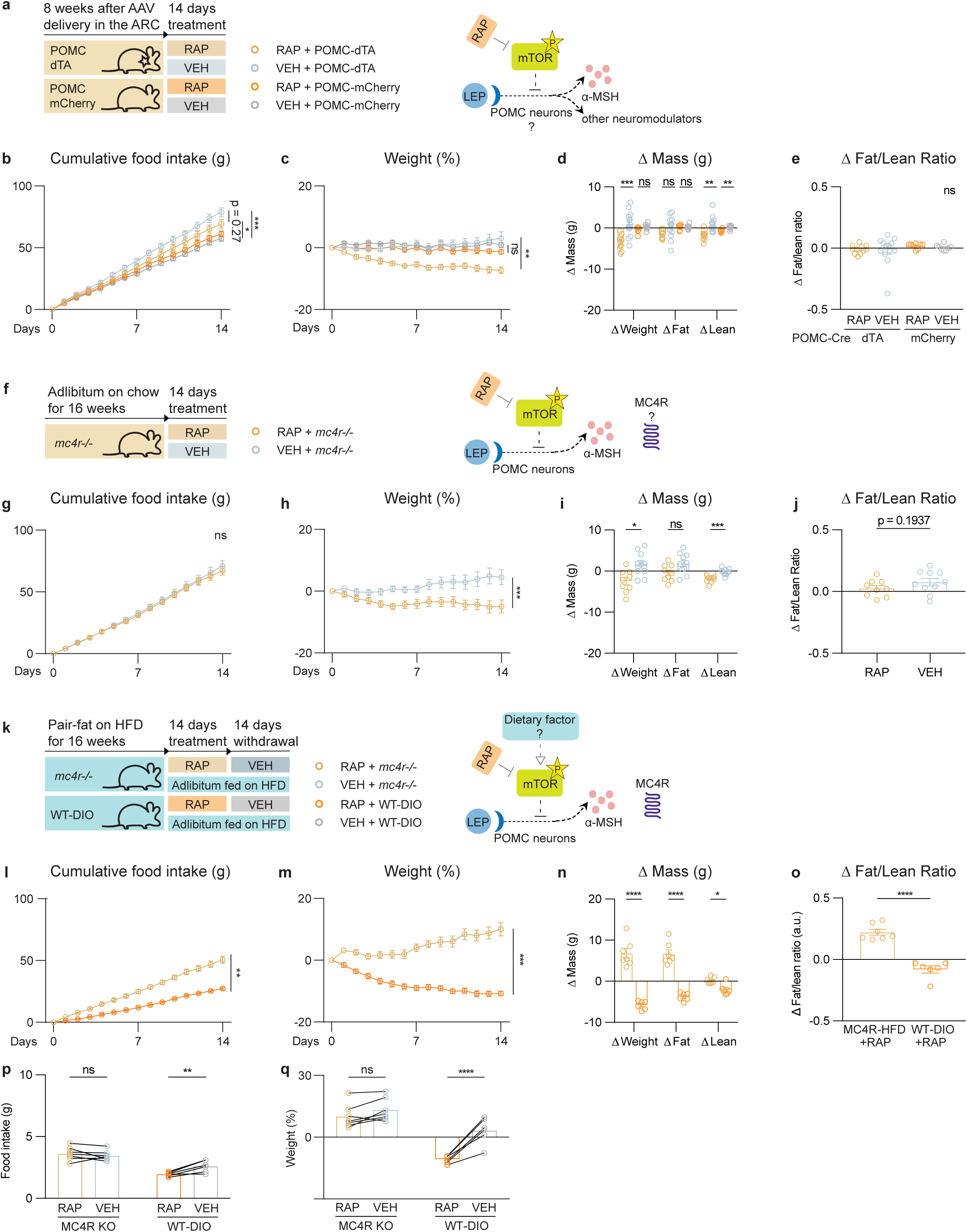
Rapamycin minimally regulates food intake, fat mass, and fat-to-lean ratio in obese mice models deficient in melanocortin signaling. (a) POMC-ablation: Schematic of POMC-dTA vs. POMC-mCherry mice that were treated with daily i.p. injections of 2 mg/kg RAP vs. VEH for 14 days. Diagram of the interaction between the leptin, mTOR, and POMC neurons. (b) Cumulative food intake, and (c) Weight, (d) Δ Mass of weight, fat and lean tissue, (e) Δ Fat/Lean Ratio (a.u.) at Day 0 vs. Day 14 (n = 10 for POMC-dTA + RAP, n=12 for POMC-dTA + VEH, n = 8 for POMC-mCherry + RAP, n = 8 for POMC-mCherry + VEH; Two-way ANOVA with Tukey’s multiple comparisons tests for Fig. 6b-d; One-way ANOVA with Tukey’s multiple comparisons for Fig. 2e). (f) MC4R knockout: Schematic of *mc4r−/−* mice fed a chow diet treated with daily i.p. injections of RAP (2 mg/kg) vs. VEH for 14 days. Diagram of the interaction between the leptin, mTOR, and POMC-MC4R pathway. (g) Cumulative food intake, (h) Weight in chow-fed *mc4r−/−* mice, (i) Δ Mass of weight, fat and lean tissue, and (j) Δ Fat/Lean Ratio (a.u.) at Day 0 vs. Day 14 (n = 8, 10, for RAP, VEH groups, respectively; Two-way ANOVA, with Šidák’s multiple comparisons for Fig. 6g-i; Two-tailed Student’s t-tests for Fig. 6j). (k) Schematic of the study design: *mc4r−/−* mice fed a HFD pairfat to WT-DIO mice for 16 weeks, after which they were treated with 14-day daily 2 mg/kg RAP i.p. injections followed by another 14-day daily i.p. VEH injections. Both groups of mice had adlibitum access to HFD throughout the pharmacological treatment. Schematic diagram of testing diet-dependent effects of rapamycin in HFD-*mc4r−/−* obese mice. Pairfat *mc4r−/−* and WT-DIO mice were treated with daily i.p. injections of RAP (2 mg/kg) for 14 days while adlibitum fed on HFD. (l) Cumulative food intake and (m) Weight, (n) Δ Mass of weight, fat and lean tissue and (o) Δ Fat/Lean Ratio (a.u.) at Day 0 and Day 14. (p) Averaged daily food intake, and (q) Weight in (previously pairfat) *mc4r−/−* vs. WT-DIO mice after withdrawal from RAP followed by VEH for 14 days (n = 7 for pairfat *mc4r−/−* mice, n = 6 for WT-DIO mice; Two-way ANOVA, with Šidák’s multiple comparisons for Fig. 6l-n, p, q; Two-tailed Student’s t-tests for Fig. 6o). All error bars represent mean ± SEM. ns, not significant, *P < 0.05, **P < 0.01, ***P < 0.001, ****P < 0.0001. Additional information is included in Fig. S8.

MC4R, a G-protein coupled receptor, is an α-MSH receptor and *mc4r−/−* mice develop obesity as do patients with mutations in this gene (Fig. S8b) ^34^. At baseline, *mc4r−/−* animals were hyperphagic and obese and these animals did not respond to leptin treatment (Fig. S8b, j, k). Similar to POMC-dTA mice, rapamycin did not reduce food intake in *mc4r−/−* mice (Fig. 6g) and while it induced a small decrease in body weight (−5.0 ± 2.1 % vs. 4.5 ± 2.4 %; p<0.001), this was attributable to loss of lean mass (RAP: –2.0 ± 0.3 g vs. VEH: –0.2 ± 0.2, n = 9, 10; p < 0.001) with no significant effect on fat mass or fat-to-lean ratio (Fig. 6h-j, S8b). A combination of rapamycin and leptin also failed to reduce weight with a significantly greater reduction of weight in the chow-fed WT controls compared to the *mc4r−/−* mutants (Fig. S8m-q). Thus despite being hyperleptinemic, in contrast to DIO mice, mice with defects in melanocortin signaling failed to respond to rapamycin.

We also controlled for possible dietary effects by feeding *mc4r−/−* mice a HFD. This was necessary because the *mc4r−/−* mice on a HFD were more obese that DIO mice. To address this, we normalized the weight of *mc4r−/−* mice to control DIO animals by feeding them 10 % fewer calories than were consumed by ad libitum DIO animals (Fig. 6k, S8c). We refer to these *mc4r−/−* animals as being ‘pair-fat’ and at 18 weeks, these and the DIO control mice weighed similar amounts. Both groups were then fed ad libitum while being treated with rapamycin for 14 days followed by another 14 days of vehicle treatment (Fig. 6k). Despite similar baseline weights, rapamycin-treated *mc4r−/−* mice consumed significantly more food than the rapamycin-treated DIO mice (*mc4r−/−*: 50.6 ± 2.9 g vs. DIO: 27.4 ± 0.9 g; p<0.01) and even gained weight during the treatment, while the rapamycin-treated DIO mice showed reduced food intake and body weight (*mc4r−/−*: +10.1 ± 2.2 % vs. DIO: –10.8 ± 0.7 %; p<0.001) (Fig. 6l, m). After 14 days, fat mass and fat-to-lean ratio were significantly higher in rapamycin-treated *mc4r−/−* mice than in the rapamycin-treated DIO mice (Fig. 6n. Δfat mass: *mc4r −/−*: 6.63 ± 1.9 g vs. DIO: –3.8 ± 0.4 g; p<0.0001; Fat-to-lean ratio: *mc4r −/−*: 0.2 ± 0.0 a.u. vs. DIO: –0.1 ± 0.0 a.u.; n = 7, 6; p<0.0001), while lean mass did not change (Fig. 6n, o). Following the cessation of rapamycin treatment, DIO mice showed a significant rebound of food intake and weight and returned to their pre-treatment levels while there was no change in the *mc4r−/−* mice (Fig. 6p, q and Fig. S9e). We next tested whether increased mTOR activity in POMC neurons is sufficient to cause leptin resistance.

### Genetic Perturbations of mTOR Activity Alter Leptin Sensitivity and Energy Balance

We examined whether mTOR activation in POMC cells can alter leptin sensitivity by breeding POMC-Cre mice to *Tsc1-flox* mice, leading to a POMC-specific deletion of the *Tsc1* gene. TSC1 encodes an endogenous mTOR inhibitor and germ line *Tsc1* knockout mice show increased mTOR activity in numerous tissues (Fig. 7a). Similar to previous reports ^35,36^, chow-fed POMC*^tsc1−/−^*mice were hyperphagic and obese (Fig. S10a-d). We then assayed leptin sensitivity in these animals by treating them with vehicle followed by leptin (Fig. 7b). In contrast to the leptin responses in the control mice (POMC*^tsc1+/-^*and POMC*^tsc1+/+^*), leptin did not reduce food intake or body weight in the POMC*^tsc1−/−^* mice (Fig. 7c, d). To control for possible independent effects of the obesity that develops in these mice at baseline, we normalized the weight of the POMC*^tsc1−/−^* mice to the control mice by pair-feeding them to chow-fed mice. The pair-fed POMC*^tsc1−/−^*mice also failed to respond to leptin and even gained weight during leptin treatment (POMC*^tsc1−/−^*: 25.3 ± 5.0 % vs. control: –2.5 ± 0.6 %; p<0.0001). At the end of leptin treatment, both pair-fed POMC*^tsc1−/−^* and control mice were treated with vehicle. The control group showed a rebound in their food consumption and weight, while the POMC*^tsc1−/−^*mice did not (Fig. 7e, f, g).

**Figure 7.**
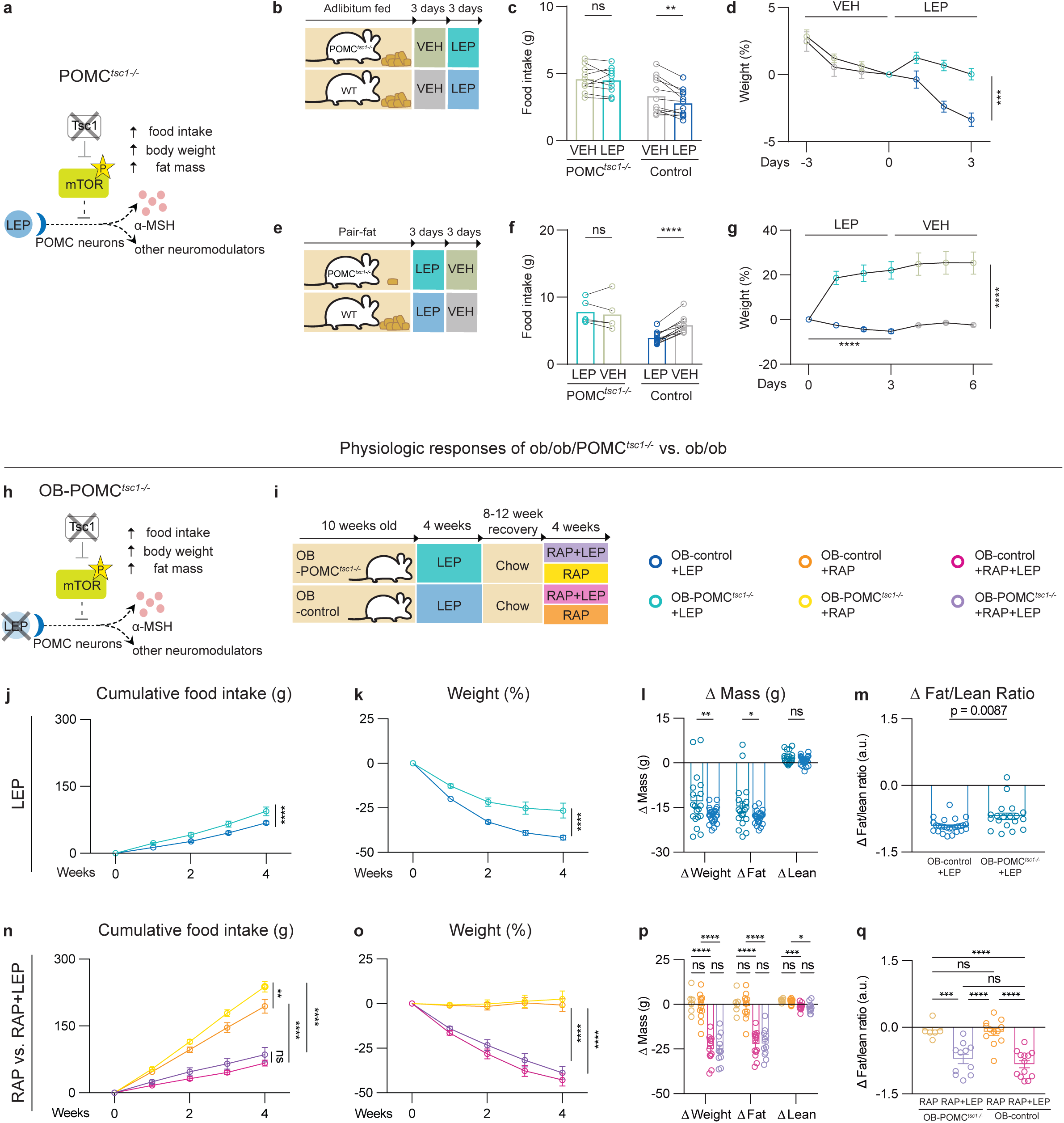
Genetic upregulation of mTOR activity in POMC cells causes leptin resistance, which is reversed by rapamycin in a leptin-signaling-dependent manner. (a) Diagram illustrates the deletion of the *tsc1* gene, resulting in mTOR activation in POMC cells and subsequent development of obesity. (b) Schematic depicting the leptin-sensitivity test in POMC*^tsc1−/−^* and WT littermate mice fed a chow diet. Both mice were treated with twice-daily i.p. injections of VEH for 3 days, followed by twice-daily i.p injections of 0.5 mg/kg LEP for another 3 days. (c) Averaged daily food intake, and (d) Weight during the 3-day leptin-sensitivity test in POMC*^tsc1−/−^*vs. WT mice ad libitum fed on chow (n = 10 for POMC*^tsc1−/−^* mice, n = 12 for WT control mice; Two-way ANOVA, with Šidák’s multiple comparisons). (e) Schematic depicting leptin-sensitivity test in pairfat POMC*^tsc1−/−^* and WT mice fed a chow diet. Both groups were treated with twice-daily i.p. injections of 0.5 mg/kg LEP for 3 days, followed by twice-daily i.p injections of VEH for another 3 days. Both groups of mice had adlibitum access to food throughout VEH and LEP treatment. (f) Averaged daily food intake, and (g) Weight during the 3-day leptin-sensitivity test in POMC*^tsc1−/−^* vs. WT mice ad libitum fed on chow (n = 5 for POMC*^tsc1−/−^* mice, n = 11 for WT control mice; Two-way ANOVA, with Šidák’s multiple comparisons). (h) Diagram illustrates a complete loss of leptin signaling in *ob/ob*/POMC*^tsc1−/−^* mice. (i) Schematic of *ob/ob*/POMC*^tsc1−/−^* (OB-POMC*^tsc1−/−^*) and *ob/ob* controls (OB-control: *ob/ob* POMC-Cre+ *tsc1fl/-* and *ob/ob* POMC-Cre-*tsc1 fl/fl*) mice fed on chow that were treated with 150 ng/hr LEP for 4 weeks, followed by an 8-to-12-week recovery period. After recovery, these mice were treated with 150 ng/hr LEP plus 2mg/kg RAP (i.p. injections 3 times a week) vs. VEH plus 2mg/kg RAP (i.p. injections 3 times a week). (j) Cumulative food intake, and (k) Weight during 4-week LEP treatment, (l) Δ Mass of weight, fat and lean tissue, and (m) Δ Fat/Lean Ratio (a.u.) at Week 0 vs. at the end of Week 4 in OB-POMC*^tsc1−/−^* vs. OB-control (n = 19, 26; respectively; Two-way ANOVA, with Šidák’s multiple comparisons for Fig. 7j-m; Two-tailed Student’s t-tests for Fig. 7m). (n) Cumulative food intake, and (o) Weight during 4-week treatment, (p) Δ Mass of weight, fat and lean tissue and (q) Δ Fat/Lean Ratio (a.u.) at Week 0 vs. at the end of Week 4 in OB-POMC*^tsc1−/−^* vs. OB-control (n = 11 for RAP + LEP in OB-POMC*^tsc1−/−^*, n = 13 for RAP+LEP in OB-control, n = 6 for RAP + VEH in OB-POMC*^tsc1−/−^*, n = 13 for RAP + VEH in OB-control; Two-way ANOVA, with Šidák’s multiple comparisons for Fig. 7n-p; Two-tailed Student’s t-tests for Fig. 7q). All error bars represent mean ± SEM. ns, not significant, *P < 0.05, **P < 0.01, ***P < 0.001, ****P < 0.0001. Additional information is included in Fig. S10.

*ob/ob* mice are leptin deficient and thus ultra-sensitive to leptin administration. We next tested whether increased mTOR activity in POMC neurons can induce complete or partial leptin resistance in *ob/ob* mice. *ob/ob* animals that carried a POMC-specific knockout of *Tsc1* were generated and at baseline, the *ob/ob*-POMC*^tsc1−/−^* double-knockout (referred to hereafter as OB-POMC*^tsc1−/−^*) mice showed a slight increase of lean mass and body weight compared to *ob/ob* controls (OB-control) (Fig. S10i, lean mass: OB-POMC*^tsc1−/−^* 22.3 ± 0.3 g vs. OB-control 19.8 ± 0.3 g; n = 26, 19; p<0.05) though the adiposity was similar between the two groups (fat mass %: OB-POMC*^tsc1−/−^* 52.3 ± 0.7 % vs. OB-control 50.9 ± 0.7 %; n = 26, 19; p = 1.0). We treated both groups with exogenous leptin for four weeks (Fig. 7i). In contrast to the OB-control mice, OB-POMC*^tsc1−/−^*showed a diminished response to leptin, characterized by 37 % greater food consumption (Fig. 7j. 94.0 ± 9.7 g vs. 68.4 ± 4.0 g; p<0.0001), 23 % higher fat mass (Fig. 7l. –14.6 ± 1.8 g vs. –19.1 ± 1.3 g, higher fat-to-lean ratio, and less weight loss than the OB-controls treated with leptin (Fig. 7k. –26.5 ± 4.2 % vs. –41.7 ± 1.3 %; p<0.0001). We next tested whether rapamycin can reverse this partial leptin resistance by treating both groups with rapamycin alone or with rapamycin plus exogenous leptin. Rapamycin alone had no effect on fat mass in the absence of leptin, while the combination of rapamycin and leptin normalized the leptin response of OB-POMC*^tsc1−/−^*mice relative to the OB-Control group with comparable reductions in food intake (Fig. 7n, RAP+LEP: OB-POMC*^tsc1−/−^*, 85.5 ± 16.2 g vs. OB-control, 66.5 ± 6.4 g; n = 13, 11; p = ns), body weight (Fig. 7o, RAP+LEP: OB-POMC*^tsc1−/−^*, –38.8 ± 3.6 % vs. OB-control, –42.8 ± 3.3 %; p = ns), fat mass (Fig. 7p, RAP+LEP: OB-POMC*^tsc1−/−^*, –22.1 ± 2.4 g vs. OB-control, –22.4 ± 2.0 g; n = 13, 11; p = ns), and fat-to-lean ratio (Fig. 7n-q, S10j). These data confirm that reduced leptin signaling in POMC neurons can cause partial leptin resistance even in *ob/ob* mice which are normally hypersensitive to leptin.

In a final set of studies, we evaluated whether mutations that blunt the activation of mTOR reduce weight gain in mice fed a HFD. Leucine and methionine activate mTOR and sensing of amino acids is mediated by the GATOR2 complex, while the RHEB GTPase directly activates the mTOR kinase and is inhibited by TSC1 (Fig. S10k) ^37–40^. POMC-specific knockouts of *Rheb* and *Wdr24*, were generated by delivering guide RNAs for these two genes, as well as control guide RNAs, into the ARC of POMC-Cas9 mice. The food intake and body weight of these groups was recorded in mice fed a chow diet for three weeks followed by a HFD for three weeks (Fig. S10l). POMC-specific knockouts of *Wdr24* or *Rheb* did not alter food intake or body weight in mice fed a chow diet (Fig. S10m, n), while both groups showed significantly reduced food intake and attenuated weight gain when fed a HFD (Fig. S10o, p). Thus, consistent with the pharmacologic effects of mTOR inhibition by rapamycin, CRISPR-mediated knockouts of these mTOR activators significantly attenuates the development of diet-induced obesity.

## DISCUSSION

Obese humans and DIO mice have increased circulating leptin levels and a diminished response to the exogenous hormone ^3,7,8^. These are the hallmarks of a hormone resistance syndrome, analogous to insulin resistance as a cause of Type 2 Diabetes ^41^, thus suggesting that obesity in DIO mice and most obese humans is caused by a ‘block’ in leptin action. The anatomic and molecular basis of this ‘block’ has been unknown and its elucidation would have important implications for our understanding of the pathogenesis of obesity as well as potential therapeutic implications. In an initial metabolomic screen for biomarkers indicative of a leptin response, we found that the levels of several mTOR ligands inversely correlate with leptin sensitivity leading us to test whether increased mTOR activity might contribute to leptin resistance. We found that rapamycin, a specific mTOR inhibitor, reduces body weight by restoring leptin signaling in DIO mice but not in mice with defects in leptin signaling or low circulating levels of the hormone. We then employed snRNA-seq to show that rapamycin treatment of DIO, but not lean mice, specifically induced genes in POMC neurons that promote leptin signaling and melanocortin production. Further studies showed that POMC neurons and melanocortin signaling are necessary for rapamycin’s weight reducing effects and that increased mTOR activity in POMC neurons is sufficient to cause leptin resistance. Thus, while the pathogenesis of insulin resistance is still not completely understood despite many decades of research, these data establish a cellular and molecular basis for leptin resistance in DIO mice. The response to treatment in DIO mice is highly predictive of a clinical response and the pathogenesis of obesity in these animals is thus considered to be equivalent to that in humans.

Our finding that the leptin resistance that develops in DIO mice is a result of mTOR activation primarily in POMC neurons is consistent with genetic studies of obesity in mice and humans ^33,34,42^. POMC encodes a protein precursor that is processed by proteases PCSK1/2 to generate α-MSH which can potently reduce food intake and body weight through broadly distributed receptors in the brain, including MC4R ^43,44^. Mutations in *Pomc*, *Pcsk1* and *Mc4r* all cause severe early onset obesity in humans ^34,42^, and setmelanotide, a peptide analogue of α-MSH, reduces weight in these individuals as well as in patients with leptin receptor mutations ^45,46^. Our data indicate that defects in melanocortin signaling can also be acquired when mice are fed a HFD. It is not yet clear why this diet leads to increased mTOR activity in POMC neurons, but a previous study has suggested that sustained hyperleptinemia can cause leptin resistance ^47–49^. We hypothesize that weight gain after consumption of a palatable diet by C57Bl/6J mice, and other strains that are obesity prone, leads to increased leptin levels in turn increasing mTOR activity as a means for blunting the response (i.e; tachyphylaxis), as has also been suggested for other hormones including insulin ^41^. Further studies will be necessary however to assess this and it is also possible that other mechanisms contribute. While an earlier report suggested that rapamycin blunted the acute effects of leptin, this study only evaluated the effect of a single dose of leptin on food intake and did not specify which neuronal type was responsible for the effect ^50^. It is thus possible that while transient mTOR activation is required for leptin’s acute effects, chronic mTOR activation, possibly induced by chronically high hormone levels, leads to a down regulation of leptin signaling.

The activity of POMC neurons is modulated by leptin, insulin, GLP1 and multiple other signals and recent studies have indicated that there is heterogeneity of POMC neurons with different clusters expressing distinct molecular markers ^24–26,28,51,52^. Our data revealed that rapamycin specifically affects the POMC subtype that expresses *Lepr*, with little effect on the other subtypes including a *Pomc+/Htr2c+* cluster. This is consistent with prior results showing that a POMC-specific deletion of *Htr2c* has only minimal effects on body weight in chow fed mice ^53^, while a deletion of *Lepr* in POMC neurons leads to increased weight ^54^.

Consistent with a role for mTOR to cause leptin resistance, our studies as well as prior studies of aged mice and mice with a deletion of *Tsc1* in POMC neurons, revealed that increased mTOR activity in these neurons causes obesity that is ameliorated by rapamycin ^35,36^. However, this previous study did not assess whether rapamycin reduced weight by restoring leptin signaling *in vivo* nor did it establish the importance of mTOR hyperactivity in POMC neurons as a cause of diet-induced obesity, the pathogenesis of which, as mentioned, is considered to be equivalent with human obesity. We also observed an improvement of POMC neural activity and leptin responsiveness after rapamycin treatment using electrophysiology. However, because DIO mice need to be fed a HFD for several months, we cannot assess the extent to which aging might also contribute to the reduced POMC neural activity of and leptin responsiveness that we observed in the slice preparations using electrophysiology.

Our results are also consistent with numerous prior reports showing that rapamycin treatment or diets low in methionine or leucine mitigate diet-induced obesity ^55–57^. However, in these prior studies, an effect on leptin sensitivity was not evaluated and the role of POMC neurons in mediating this response was not shown. Indeed, we find that downregulation of the leucine-sensing mTORC1 pathway in POMC neurons blunts weight gain in POMC-*Wdr24* knockout mice fed a HFD. Thus overall, our data unify and extend a large number of prior studies by showing that increased mTOR signaling in POMC neurons is both necessary and sufficient for the development of diet-induced obesity and provides a mechanism explaining how and why low protein diets and rapamycin have been previously shown to reduce obesity.

We also found that increased mTOR activity in POMC neurons of *ob/ob* (by breeding to POMC*^tsc1−/−^* mice) leads to partial leptin resistance in these animals, which are normally extremely sensitive to leptin. However, leptin’s effect, while reduced, is still significant in these mice suggesting that the hormone acts on other target populations including AgRP neurons, an arcuate population that drives appetite and is inhibited by leptin, and likely others. Our finding that DIO is caused by defects primarily in POMC neurons is consistent with the fact that *ob/ob* mice are significantly more obese than DIO mice, likely as a result of leptin action at these other cellular sites besides POMC neurons. Thus our data do not exclude the possibility that mTOR might also affect leptin signaling in additional cell types. Consistent with this, we observed increased pS6 levels in a number of hypothalamic sites outside the ARC in DIO vs. chow fed mice such as the ventromedial hypothalamus, preoptic areas as well as in the basomedial amygdala and striatum. It is not clear which of these neurons express the leptin receptor and what their possible contributions to leptin resistance are.

The gene expression data suggest that chronic mTOR activation activates genes that diminish leptin signaling and that rapamycin reverses this. Our finding that rapamycin increases the levels of pSTAT3 provides one of several potential mechanisms by which leptin signaling is restored including decreasing the expression of *Ptprm* and *Ptprt*, which dephosphorylate pSTAT3. However other mechanisms in addition to dephosphorylation of pSTAT3 may also contribute because a POMC-specific knockout of *Stat3* results in only a mild obese phenotype ^5,58^. Other signaling molecules have been shown to contribute to leptin resistance including SOC3 and PIAS proteins that inhibit JAK and STAT activity respectively to decrease leptin signal transduction ^58–65^. Another report suggested that increased mTOR signaling in POMC neurons increases the activity of an inhibitory K^ATP^ channel and that POMC neurons from POMC*^tsc1−/−^* mice are hyperpolarized via increased K^ATP^ conductance ^35^. Other reports also show that increased levels of PIP3 can silence POMC neurons by increasing K^ATP^ channel activity ^66^. Thus increased expression of the K^ATP^ channel or similar channels could also reduce the activity of POMC subtype 2 neurons expressing the leptin receptor. Recent reports have suggested that a knockout of *Grb10*, an endogenous mTOR inhibitor, in POMC neurons enhances leptin signaling ^4,67^ but, similar to the minimal effects of a POMC knockout of *Lepr* and *Stat3*, a POMC-specific knockout of this gene did not alter the weight of chow-fed mice to the same extent as did a knockout of *Tsc1* in our and the aforementioned studies ^35,36^. Inhibiting HDAC6 has also been reported to alleviate leptin resistance though in this case the effect appeared to be peripheral, possibly leading to alterations of signaling onto POMC neurons to alter mTOR activity ^68^. Additional reports have suggested that impaired calcium influx, autophagy and/or increased ER stress in POMC neurons can cause leptin resistance ^69–71^, though the metabolic impairment seen in mice with mutations in *Atg7* in POMC neurons did not lead to the same level of obesity that develops in DIO mice or mice with a POMC deletion of *Tsc1* ^70^. Still other studies have suggested that ER stress contributes to leptin resistance which can be alleviated by the drug celastrol and while a possible role for the IL1 receptor was shown ^72–76^, its precise cellular target is unknown. Finally, several GI hormones including CCK, amylin and dual and triple agonists can also restore leptin action in DIO mice but the neural mechanism is as yet unknown ^77,78^. Further studies will thus be necessary to integrate these findings and fully establish whether these agents alter mTOR signaling in POMC and other neurons or whether other signaling pathways can also regulate leptin sensitivity.

Rapamycin was first developed clinically as an immune suppressor ^79,80^. More recently it is being evaluated as a possible agent for extending lifespan but the mechanism responsible for the potential effects on longevity are not fully understood ^81^. Increased BMI has been shown to be an independent risk factor for mortality ^82^, raising the possibility that restored leptin sensitivity and reduced body weight could contribute to rapamycin’s effect on longevity. However, rapamycin also leads to glucose intolerance and insulin resistance which would generally limit its utility, especially in obese patients with prediabetes and diabetes ^19,79,80^. We also found that, while not worsening diabetes, rapamycin treatment of DIO mice has only a marginal effect to improve glucose metabolism in DIO mice. It seems likely that the benefit of weight loss is counteracted by the negative effect of rapamycin on insulin signaling. Consistent with this, we found that co-administration of rapamycin and leptin significantly ameliorated glucose intolerance in wild-type mice but not in mice with defects in melanocortin signaling. This is consistent with prior studies showing that leptin further improves glucose tolerance in mice with mutations in *Akt* and that POMC neurons have effects on glucose metabolism independent of their effects on body weight^23,70,83–85^.

While rapamycin specifically reduced fat mass in animals with intact leptin signaling, we also observed highly reproducible effects of rapamycin to decrease lean mass including animals with defects in leptin and melanocortin signaling. The mechanism accounting for this effect is unknown though prior reports have shown that rapamycin can decrease protein synthesis ^14,15^. We also found that rapamycin elicited a modest effect to increase energy expenditure in DIO mice. However the effects on lean mass in DIO mice with functional leptin signaling were minimal and significantly smaller than the effect on fat mass, consistent with numerous prior reports showing that leptin specifically reduces fat mass with little or no effect on lean mass (in leptin-sensitive animals) ^2,3^. Thus, the effect of rapamycin on lean mass is likely to be independent of leptin possibly via the aforementioned inhibition of physiological protein synthesis ^14,15^. Prior reports have shown that an adipose-tissue knockout of *Raptor*, a component of mTORC1, prevents weight gain of animals fed on HFD ^86^, while an adipose-tissue knockout of *Rictor*, a component of mTORC2, exacerbates weight gain on HFD ^87^. However, these studies did not evaluate effects on lean mass, and it is possible that rapamycin might also have other effects on other tissues.

Overall, these data suggest that selective inhibition of mTOR in POMC neurons could provide a new strategy for leptin re-sensitization as a treatment for obesity. While several new incretin-based therapies have shown potent effects on reversing obesity, it is likely that other therapeutic approaches will be necessary for managing obesity in patients who cannot tolerate these drugs, fail to respond to them or to help maintain weight in patients after treatment ^88,89^. There is also significant recidivism after weight loss post bariatric surgery which further highlights the need to develop additional means for weight maintenance ^90^. Leptin levels decrease after weight loss and some of the weight gain that is seen after bariatric surgery or dieting appears to be a result of this ^91^. Thus, new therapies that enhance endogenous leptin sensitivity via cell-type-specific inhibition of mTOR activity may provide an important therapeutic modality on its own or as an adjunct to incretin therapies or bariatric surgery. Recent advances in developing brain-specific rapalogues provide a possible means to selectively reduce mTOR activity specifically in brain and thus represent a potential means for treating of brain diseases associated with mTOR hyperactivity ^92–95^. Alternatively, the development of means for cell specific delivery of rapamycin or identifying cell-type-specific mTOR components in POMC neurons could provide new avenues for treating obesity or maintaining weight loss ^96^.

In summary, we show that leptin resistance in DIO animals is caused by increased mTOR activity in POMC neurons, and that rapamycin reduces obesity by re-sensitizing endogenous leptin signaling in these cells. These findings thus have important implications for our understanding of the pathogenesis of obesity and potential therapeutic applications.

## AUTHOR CONTRIBUTIONS

B.T. conceived and led the study, designed and conducted the experiments, analyzed the data, conceptualized the results, wrote the manuscript and provided the funding support. K.H. designed and conducted the experiments, bred mouse models, wrote the manuscript and conceptualized the results. L.K. conducted electrophysiological recordings and analysis. J.D.L. conducted quality control and initial analysis of single-nucleus RNAseq datasets. Z.Z. conducted metabolomic and lipidomic sample preparation and analysis. J.D.R. provided resources for metabolomics and lipidomics. J.M.F. conceived and led the study, wrote the manuscript and conceptualized the results, provided resources and funding support.

## ACKNOWLEDGEMENTS

We thank Cori Bargmann and Nat Heintz for their advice throughout the project and their critical comments on the manuscript. We thank Donghoon Lee for the help with tissue dissections. We thank Alexandre Moura Assis for their help with feeding animals and proofreading the manuscript. We thank Lisa Pomeranz, Max Halaas for feeding animals. We thank Hector Lugo for care of animals. We thank Xianfeng Zeng and Wenyun Lu for assisting Z.Z. to perform sample preparation, lipidomics analysis and develop the LC methods. We thank Connie Zhao and Hong Duan for their help with 10X genomics library preparation and sequencing. B.T. acknowledges support from the David Rockefeller Fellowship. J.M.F. acknowledges support from the JPB foundation. B.T. and K.H. acknowledge support from the Robertson Therapeutic Development Fund, and the clinical and translational science awards.

## METHODS

### Animals

Wild-type mice (#000664), *db/db* mice (#000697), *mc4r−/−* (#032518), POMC-Cre (#005965) *Tsc1 fl/fl* (#005680), AgRP-Cre (#012899), LSL-Cas9 (#026175), POMC-EGFP (#009593) were acquired from Jackson Lab and *ob/ob* were F1, bred in lab, from a cross between male *ob/ob* and female *ob/+* (#000632). All crosses were bred in lab from the above animals. Animals were kept at ambient temperature and humidity-controlled housing with a 12hr light-dark cycle (lights on at 7am and off at 7pm) and on a standard-chow diet (PicoLab® Rodent Diet 205053) unless otherwise indicated. Diet-induced-obese (DIO) wild-type mice were fed on high-fat diet (HFD, Research Diets, Cat# D12492, Rodent Diet With 60 kcal% Fat) starting at ∼6 weeks old and used for experiments starting at 24 weeks old (fed on HFD for at least 18 weeks). Aged wild-type mice (∼15 months old, #000664) were acquired from Jackson Lab. All experiments were conducted according to AAALAC approved animal protocols #18050, #18051, #22012 and #21064. Males were used throughout. Males and female animals were used in Figure 2g-2f, 3a-d, and 7h-q. All experiments were internally sex and age matched.

### Plasma metabolomics and lipidomics

Wildtype animals (Jackson #000664) were fed either chow (PicoLab® Rodent Diet 205053) or high fat diet (HFD, Research Diets cat. #D12492 Rodent Diet With 60 kcal% Fat) starting at 6-9 weeks of age. *ob/ob* males, bred in lab as described above, were also fed either chow or HFD and pairfed to their wildtype counterparts. Food intake for the wildtype animals were measured weekly, divided by 7, and then fed in that amount daily to the pairfed *ob/ob* animals for 18 weeks. Despite this pairfeeding scheme, the *ob/ob* animals gained more weight than their wild-type counterparts. After 18 weeks of daily pairfeeding, all animals were i.p. injected with PBS (Gibco, Cat# 14190-144) at 12 hour intervals for 24hrs and blood was collected and processed as described below in section (7). Following 5 days of recovery, the animals were i.p. injected with 12.5 mg/kg leptin, dissolved in PBS at 12 hour intervals for 24hrs and blood was again collected. (1) Metabolomics. For polar metabolites, 10ul of serum was extracted with 40ul MeOH pre-cooled on dry ice. After vertexing, samples are left on dry ice for 5min then centrifuged at 16,000 X g for 10 min at 4°C. Supernatant was subjected to LC-MS analysis. Positive and Negative mode of metabolomics were run on a quadrupole-orbitrap mass spectrometer (Q Exactive, Thermo Fisher Scientific, San Jose, CA) coupled with hydrophilic interaction chromatography (HILIC) via electrospray ionization. LC separation was done with a XBridge BEH Amide column (2.1 mm x 150 mm x 2.5 mm particle size, 130 A ° pore size; Waters, Milford, MA) using a gradient of solvent A (20 mM ammonium acetate, 20 mM ammounium hydroxide in95:5 water:acetonitrile, pH 9.45) and solvent B (acetonitrile). Flow rate was 150 mL/min. The LC gradient was: 0 min, 85% B;2 min, 85% B; 3 min, 80% B; 5 min, 80% B; 6 min, 75% B; 7 min, 75% B; 8 min, 70% B; 9 min, 70% B; 10 min, 50% B; 12 min,50% B; 13 min, 25% B; 16 min, 25% B; 18 min, 0% B; 23 min, 0% B; 24 min, 85% B. Injection volume was 5ul for all serum samples at the autosampler temperature of 5 °C. (2) Lipidomics. Serum lipidomic samples are extracted with ethyl acetate. Serum (4μl) was added to ethyl acetate (100μl) and centrifuged 16,000 X g for 10min, and the supernatant was collected. The same process was repeated, and supernatant is combined. The resulting extract was dried down and redissolved in 1:1:1 methanol:acetonitrile:2-propanol (200μl) before analysis by Q Exactive Plus mass spectrometer coupled to a Vanquish UHPLC system (Thermo Fisher Scientific) using positive and negative-mode electrospray ionization. The LC separation was achieved on an Agilent Poroshell 120 EC-C18 column (150 × 2.1mm, 2.7 µm particle size) at a flow rate of 150 µl min^−1^. The gradient was 0minutes, 25% B; 2minutes, 25% B; 4minutes, 65% B; 16minutes, 100% B; 20minutes, 100% B; 21minutes, 25% B; 27minutes, 25% B. Solvent A is 1mM ammonium acetate + 0.2% acetic acid in water:methanol (90:10). Solvent B is 1mM ammonium acetate + 0.2% acetic acid in methanol:2-propanol (2:98).

### Pharmacological administration

Recombinant mouse leptin (R&D 498-OB-05M) was dissolved in PBS and injected intraperitoneally (i.p.) or delivered via a subcutaneous osmotic pump (Alzet Cat# 2002, 2004, or 2006). Osmotic pumps were filled and calibrated using the manufacturer’s instructions. They were inserted dorsally under the skin of an isoflurane-anesthetized mouse using sterile surgery techniques. Rapamycin (LC Laboratories, Cat# 53123-88-9) was first dissolved in DMSO at 200 mg/ml, then diluted in 5% PEG 400 and 5% Tween 80 (in PBS) to a final concentration of 0.5 mg/ml. I.p. injections were done at indicated concentrations using insulin syringes (Beckton Dickinson, Cat# 324911).

### Magnetic resonance imaging (MRI)

Body fat mass was measured by MRI using Echo-MRI 100H (EchoMRI, LL). Body fat percentage was calculated by dividing fat mass over total body mass. Lean mass was calculated by subtracting fat mass from total body mass.

### Leptin sensitivity test

Animals were administered with leptin (at indicated doses) or PBS either by i.p. injections every 12 hours or via osmotic pumps implanted 1 day prior to the start of the experiment. Food intake and body weight were measured through the course of treatment as indicated.

### Glucose tolerance test

Animals fasted overnight were i.p. injected 5-20% glucose dissolved in PBS as indicated. Total amount of glucose injected was based on lean mass times the dosage. Dosage used for each experiment and cohort was indicated in the manuscript. Blood glucose in the GTT assay and ad libitum-fed conditions were measured by tail vein sampling using a Breeze2 glucometer (Bayer SKU: breeze2meter UPC: 301931440010). For the GTT in DIO mice, the groups of mice receiving 600 ng/hr leptin were i.p. injected 1 mg/kg leptin 1 hour prior to the glucose injection, while the other groups of mice were i.p. injected PBS 1 hour prior to the GTT started.

### Leptin ELISA

Blood was collected retro-orbitally using EDTA coated capillaries (Drummond Calibrated Micropipettes Glass Capillaries with EDTA 100µl, Cat# 2-000-100-D). Samples were spun for 20 minutes at 4°C and supernatant collected as plasma and frozen immediately in liquid nitrogen and stored at –80°C in screw cap tubes. Plasma leptin was measured by ELISA (Alpco, Cat# 22-LEPMS-E01) according to the manufacturer’s protocol.

### Pairfeeding or Pairfat conditions

Pairfeeding was conducted by measuring the daily or weekly food intake of the group which shows lower food intake and feeding that same amount to the other group. In some conditions where the group being pairfed still weighs significantly higher than the group it is pairfed to, we further restricted their food intake 10% lower than the amount of food they received in order to reach equal body weight (Pairfat) as the other group before the subsequent tests. Animals were single housed during experiments. Food intake and body weight were measured daily or weekly using an Ohaus Scale. For pairfeeding in Figure 1, pairfeeding was done daily; however the amount fed was determined but measuring the food intake of the WTs once a week and dividing that by 7 and then feeding the OB animals that amount daily in the following week. For Figure 3e-3h, during weeks 19-23 (21 days), vehicle treated animals were fed, daily, what the rapamycin treated animals had eaten the preceeding 24hours. For Figure 6k, animals were pairfed on HFD, food amount determined weekly in the first 16 weeks with the *mc4r−/−* cages receiving 10% less food than consumed by the wildtype animals. The last 3 weeks, animals were single housed and the *mc4r−/−* animals were precisely pairfed individually on a daily basis 10% less than what the wildtype animals were eating. This resulted in all groups weighing the same at the beginning of the ensuing leptin sensitivity tests. We refer to this protocol as “Pairfat”.

### Whole-brain mTOR activity (pS6) mapping

SHIELD-based whole-brain clearing and labeling was employed for mapping pS6 activity in DIO mice and chow fed mice. Mice ad libitum fed on HFD or chow were anesthetized with isoflurane and transcardially perfused with PBS containing 10 U/ml heparin, followed by 4% PFA. The dissected brains were fixed in 4% PFA for 24 h at 4 °C. Brains were then transferred to PBS containing 0.1% sodium azide until brain clearing and labeling. Brains were processed by LifeCanvas Technologies following the SHIELD protocol as previously published ^21,97^. Samples were cleared for 7 days with Clear+ delipidation buffer, followed by batch labeling in SmartBatch+ with and 5 μg anti-Rabbit pS6 (Invitrogen Cat# 44-923G) per brain. Fluorescently conjugated secondary antibodies were applied in 1:2 primary/secondary molar ratios (Jackson ImmunoResearch). Labeled samples were incubated in EasyIndex (LifeCanvas Technologies) for refractive index matching (n = 1.52) and imaged with SmartSPIM (LifeCanvas Technologies) at 4 μm z-step and 1.8 μm xy pixel size. Image analysis was conducted following the procedures as previously published ^21,97^.

### Single-nucleus RNA sequencing

Animals received 3-day treatment of rapamycin and/or leptin were anesthetized under 5% isoflurane 4-hour post last injection. ARC was microdissected under a stereo microscope in pre-chilled dissection buffer and immediately transferred to dry ice prior to downstream nuclei extraction at the same day. After dissection, frozen tissues on dry ice were immediately transferred to Teflon homogenizer containing 1 ml pre-chilled NP40 lysis buffer (Fisher Cat# FNN0021) and homogenized for 15-30 times using a pellet pestle on ice. Homogenized samples were incubated for another 10-15 min on ice, followed by passing through a 70 µm Flowmi Cell Strainer and a 40 µm Flowmi Cell Strainer (Millipore Sigma). The collected flowthrough was centrifuged at 500-1000 rcf for 5 min at 4°C and pellets were resuspended completely in 1 ml 20% iodixanol at 4°C. Samples were centrifuged at 10000 rcf for 20 min at 4°C. Supernatant containing debris was carefully removed. Pellets were resuspended in staining buffer containing anti-NeuN Alexa 647 antibody (abcam, Cat# ab190565) and HashTag antibodies (1 µl for labeling ∼1 x 10^6^ nuclei, BioLegend, TotalSeqB anti-Nuclear Pore) in order to enrich neurons and multiplex samples. After antibody incubation and rotating for 30 min at 4°C, samples were washed with staining buffer without antibodies for 3 times. Samples were resuspended in FACS buffer after last-round wash and sent for FACS sorting. Hoechst 33342 (ThermoFisher Scientific, Cat# H3570) were added at a final concentration of 0.2 mM to label nuclei. Sorted nuclei were sent for downstream 10X genomics 3’ RNA-seq with feature barcode library preparation and sequenced using NovaSeq sequencer or DNBseq sequencer at BGI with ∼30000 reads/nuclei on average. Dissection buffer contains 1X HBSS, 2.5 mM HEPES-KOH [pH 7.4], 35 mM Glucose, 4 mM NaHCO_3_, and Actinomycin D (Sigma-Aldrich, Cat# A1410) at a final concentration of 20 µg/ml. NP40 lysis buffer contains 10 mM Tris-HCl [pH 7.4], 10 mM NaCl, 3 mM MgCl_2_, 0.1% NP40 dissolved in nuclease-free water. For 1 ml NP40 lysis buffer, 1 µl DTT, 25 µl 20 U/µl SupeRasine (Thermo Cat# AM2696), 12.5 µl 40 U/ µl RNasin (Promega Cat# N2615), 10 µl protease and phosphatase inhibitor cocktail (100X; Thermo Cat# 78442), 40 µl 1 mg/ml Actinomycin D were added right before use. 20% iodixanol buffer contains 0.25M sucrose, 25mM KCl, 5mM MgCl_2_, 20mM Tricine-HCl [pH 8.0] and 20% Iodixanol dissolved in nuclease-free water. DTT, Superasine, Rnasin and protease inhibitors were added at the same concentration as NP40 lysis buffer right before use. Staining buffer contains 2% BSA, 0.05% NP40 dissolved in nuclease-free 1X PBS. Superasine, Rnasin and protease inhibitors were added at the same concentration as NP40 lysis buffer right before use. FACS buffer contains 2% BSA dissolved in nuclease-free 1X PBS buffer. Superasine, Rnasin and protease inhibitors were added at the same concentration as NP40 lysis buffer right before use.

### snRNA-seq analysis

The fastq files were aligned to mouse genome (mm10), and the expression levels in each cell were estimated with Cellranger (v 6.0.0). The gene expression count matrix for each sample was processed with the following steps: (1) Estimate doublet with Scrublet (https://github.com/swolock/scrublet) ^98^; (2) Estimate and correct the ambient RNA contaminations with SoupX (https://github.com/constantAmateur/SoupX) ^99^; (3) Load the corrected counting matrix into Seurat object with log normalization; (4) Calculate the proportion of UMIs from mitochondrial genes; (5) Demultiplex with hashtag oligos followed the Seurat vignette (https://satijalab.org/seurat/articles/hashing_vignette.html); (6) The cells assigned as doublets or mitochondrial content greater than 1% were removed. The Seurat objects were integrated by following the RPCA workflow (https://satijalab.org/seurat/articles/integration_rpca.html) ^100^. The number of PCs used for UMAP calculation was selected with elbow plot ^101^. Then, the clustering was calculated with Leiden algorithm ^102^. To select the optimized resolution, the resolution was tested from 0.1 to 1.0 and was selected with clustree. The Chow_HFD was assigned as the reference dataset. The clustering information were mapped and transferred from the reference to OB_RAP_VEH datasets by following the Seurat vignette (https://satijalab.org/seurat/articles/integration_mapping.html).

The differential gene expression of each comparison was performed with Seurat::FindMarkers() with logfc.threshold greater than 0.14, and raw p values were corrected using p.adjust() with the ‘BH’ method. Significant differential expression genes were defined as log2 (Fold Change) greater than 0.26 or less than –0.26, and corrected p values less than 0.05.

### Histology

Mice were transcardially perfused with PBS (Fisher, Cat# BP39920) followed by 4% paraformaldehyde (PFA, EMS Cat# 15714-S). Brains were dissected and post-fixed in 4% PFA at 4°C overnight. Brains were sectioned into 50-μm coronal slices using a vibratome (Leica). For immunohistochemistry, brain sections were blocked (0.1% Triton X-100 (Thermo Cat# 85111) in PBS, 3% bovine serum albumin (Sigma Cat# A9647-500G), 2% normal donkey serum (Jackson ImmunoResearch Cat# 017-000-121)). For pSTAT3 staining, sections were first rinsed in 1% H2O2 + 1% NaOH in H2O for 20 min, at room temperature, then transferred to 0.3% glycine in 1X PBS for 10 min, at room temperature prior to blocking. Sections post blocking were then incubated with primary antibody (rabbit anti-Phospho-S6, Invitrogen, Cat# 44-923G, 1:1000 dilution; rabbit anti Phosphp-Stat3, Cell Signaling, Cat# 9145 1:500 dilution; chicken anti GFP, 1:1000 dilution, abcam, Cat# ab13970) for 2 days at 4°C. Sections were then washed and incubated with secondary antibody (donkey anti-chicken IgG Alexa 488, Jackson Immunoresearch, Cat# 703-546-155, 1:1000 dilution; donkey anti-rabbit IgG Alexa 594, Invitrogen, Cat# A21207, 1:500 or 1:1000 dilution) for 1 hour at room temperature, washed again, mounted with DAPI Fluoromount-G (Southern Biotech Cat# 0100-20) and imaged with SlideView microscope (VS200, Olympus). Images underwent minimal processing (such as adjusting brightness and contrast) performed using ImageJ. The CellCounter plugin for ImageJ was used to quantify numbers and percentages of co-localizations from sections.

### Slice preparation and electrophysiology

Acute coronal hypothalamic brain slices (200 μm) were prepared from POMC-eGFP (8-44 weeks old). Mice fed either on chow diet, or high fat diet for a minimum of 16 weeks. Chow data were generated from 8-12 week old mice (n=7, 3-8 neurons per mouse). DIO data were generated from 20-44 week old mice (n=33, 1-10 neurons per mouse). Vehicle-treated DIO data were generated from 33 week old mice (n=4, 6-12 neurons per mouse) and Rapamycin-treated DIO data were generated from 32-33 week old, (n=4, 5-10 neurons per mouse). Mice were anesthetized with isoflurane prior to decapitation and removal of the entire brain which was immediately submerged in ice-cold ‘slicing’ solution containing (in mM): 85 NaCl, 2.5 KCl, 0.5 CaCl2, 4 MgCl2, 25 NaHCO3, 1.25 NaH2PO4, 64 sucrose, 25 glucose. This solution was bubbled with 95% O2 and 5% CO2, pH 7.4. Coronal hypothalamic slices (200μm) were made with a moving blade microtome (VT1000S, Leica). The slices were kept at 32 °C for 40 min in recording solution containing (in mM) 125 NaCl, 2.5 KCl, 1.25 NaH2PO4, 26 NaHCO3, 10 glucose, 2 CaCl2 and 1 MgCl2, pH 7.4 when bubbled with 95% O2 and 5% CO2 before being kept at room temperature prior to recording. Cell-attached patch-clamp recordings were made in voltage clamp configuration with 0pA holding current at room temperature from eGFP-expressing neurons in the arcuate nucleus. Neurons were visualized using epifluorescence and patched under DIC imaging on an upright microscope (Zeiss Axioskop 2FS Plus) equipped with a CCD camera (Hamamatsu). Patch pipettes pulled (Narishige PP-830) from borosilicate glass (Sutter Instrument) had tip resistances of 7–11 MΩ and were filled with K-gluconate internal containing (in mM): 135 potassium gluconate, 4 KCl, 0.05 EGTA, 10 Hepes, 4 MgATP, 10 Na-Phosphocreatine, pH adjusted to 7.3 with KOH, 290 OSM. All chemicals were obtained from Sigma. Leptin stock was prepared in PBS (1mg/ml or 100µM), then diluted in ASCF (100nM) and bath applied for 20 minutes via perfusion. Recordings were acquired with an Axopatch 200B amplifier, filtered to 2 kHz and digitized at 10 kHz (pClamp10 software, Molecular Devices). Data were analyzed using IGOR Pro (Wavemetrics) and NeuroMatic (http://www.neuromatic.thinkrandom.com/). Mean firing rate was detected in 10 second bins and maximum firing rate was recorded in each condition. Baseline firing rate was calculated in the first minute of recording, ASCF was calculated in the minute before peptide application. Neurons were considered excited or inhibited by leptin if firing changed more than 10%. For i.p. injections DIO mice fed on high fat diet for more than 16 weeks were injected either with rapamycin (2 mg/kg) or vehicle once a day for 3 consecutive days before brain slice preparation.

### Viral injections into the ARC

Cre-inducible AAV5-hsyn-DIO-mCherry (Addgene Cat# 50459-AAV5) and AAV5-mCherry-flex-dtA (plasmid was obtained from Addgene Cat# 58536, AAV viruses were packaged at Janelia Research Campus) viruses were injected into ARC of POMC-Cre+ animals using stereotactic surgery as follows: Mice were anesthetized with 3% isoflurane in oxygen, placed in a stereotaxic apparatus (Kopf Instruments) and kept at 1.5% isoflurane during surgery. Viruses were bilaterally delivered using a borosilicate glass pipette connected to Nanoject (Drummond, Cat# 3-000-203-G/X) in ARC (200 nl per coordinate, 50 nl/min). The coordinates to locate ARC were relative to the bregma: AP/DV/ML = −1.75/−5.95/±0.25 mm, AP/DV/ML = −1.75/−5.75/±0.25 mm. After finishing injection at each coordinate, syringes were held at the coordinate for another 3 minutes. Animals were allowed to recover (and adjust eating habits) for 8 weeks post-surgery, before being used in experiments. Plasmids containing guide-RNAs were designed, customized and generated at VectorBuilder. Plasmids were then packaged into AAV viruses at Janelia Research Campus.

### Metabolic cages

Diet-induced obese mice were singly housed in a climate-controlled (temperature: 22 °C; humidity: 55%; 12-hour light–12-hour dark cycle) automated home cage phenotyping system (TSE) with ad libitum access to water and high-fat-diet. After 7 days of adaptation to the TSE cage, receiving daily mock IP injections, the mice were treated daily with IP vehicle or IP 2 mg/kg Rapamycin for 6 days (Fig. S4). Data were collected and analyzed as recommended by the manufacturers. Energy expenditure and respiratory exchange ratio (RER) were measured by an indirect gas calorimetry module recording open circuit oxygen consumption and carbon dioxide production (VO_2_ and VCO_2_). Locomotor activity was recorded as beam breaks converted into distance/velocity, measuring activity in three dimensions and analyzed in the metabolic cage using custom software. Statistical assessment of the data was conducted using CalR software (https://www.calrapp.org).

### Statistics and reproducibility

We conducted statistical analyses using GraphPad Prism 9.0. Throughout the paper, values were reported as mean ± SEM (error bar). Each statistical test performed was denoted in the figure legends. P-values for pair-wise comparisons were obtained using two-tailed student’s t-test. P-values for two independent group comparisons were either obtained using two-tailed student’s t-test or nonparametric Mann-Whitney test based on data distributions. P-values for multiple group comparisons were conducted using one-way or two-way ANOVA (with repeated measures when possible) based on the number of factors and corrected for multiple comparisons using Fisher’s LSD test, Tukey’s test or Sidak’s test, indicated in the figure legends. The experiments were not randomized. Data for each experiment were repeated with at least two different cohorts. Representative trial data or pooled data from multiple cohorts were used for conducting statistics and plotting figures. The investigators were not blinded to allocation during experiments and outcome assessments.

## SUPPLEMENTARY FIGURE LEGENDS

**Figure S1.**
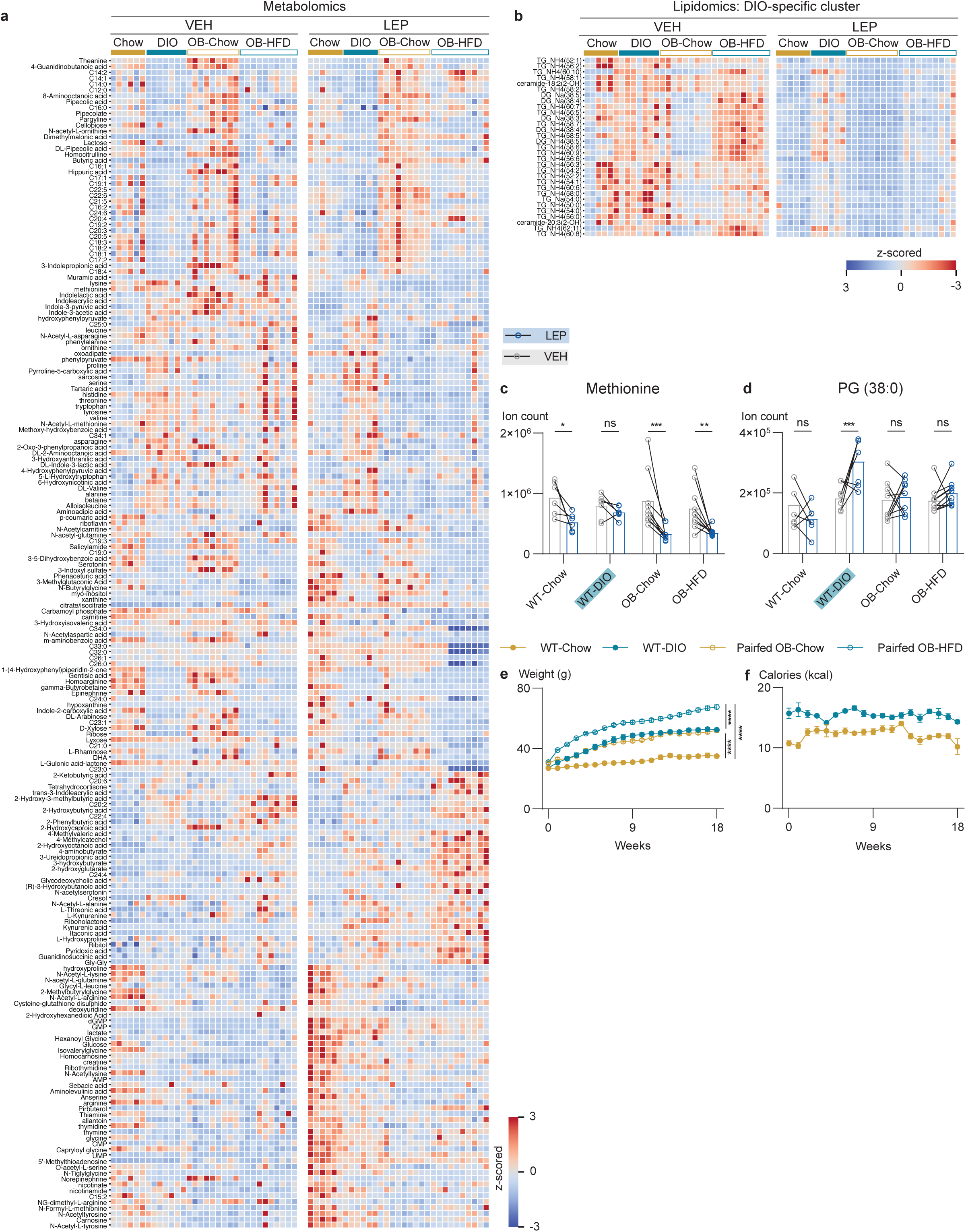
Overview of metabolomic and lipidomic plasma profiles in response to vehicle or leptin across Chow, DIO, OB-Chow and OB-HFD mouse models. (a) Heatmap overview of metabolomics and (b) DIO-cluster specific lipidomics in WT-Chow, WT-DIO, OB-Chow, and OB-HFD treated with VEH vs. LEP. Comparisons of representative metabolite levels (ion count) post VEH vs. LEP: (c) Methionine, (d) PG (38:0) (Two-way ANOVA, with Fisher’s LSD comparisons, n = 6 for WT-Chow, n = 6, 7 for DIO treated with VEH vs. LEP, n = 9 for OB-Chow, n = 10 for OB-HFD). (e) Weight during 18-week pairfeeding scheme (Two-way ANOVA, with Tukey’s multiple comparisons, n = 6, 6, 9, 10 for WT-Chow, WT-DIO, OB-Chow, OB-HFD, respectively). (f) Weekly consumption of calories (kcal) during 18-week pairfeeding scheme in WT-Chow vs. WT-DIO mice. (Two-way ANOVA, Šidák’s multiple comparisons, n = 6 for each group). All error bars represent mean ± SEM. ns, not significant, *P < 0.05, **P < 0.01, ***P < 0.001, ****P < 0.0001.

**Figure S2.**
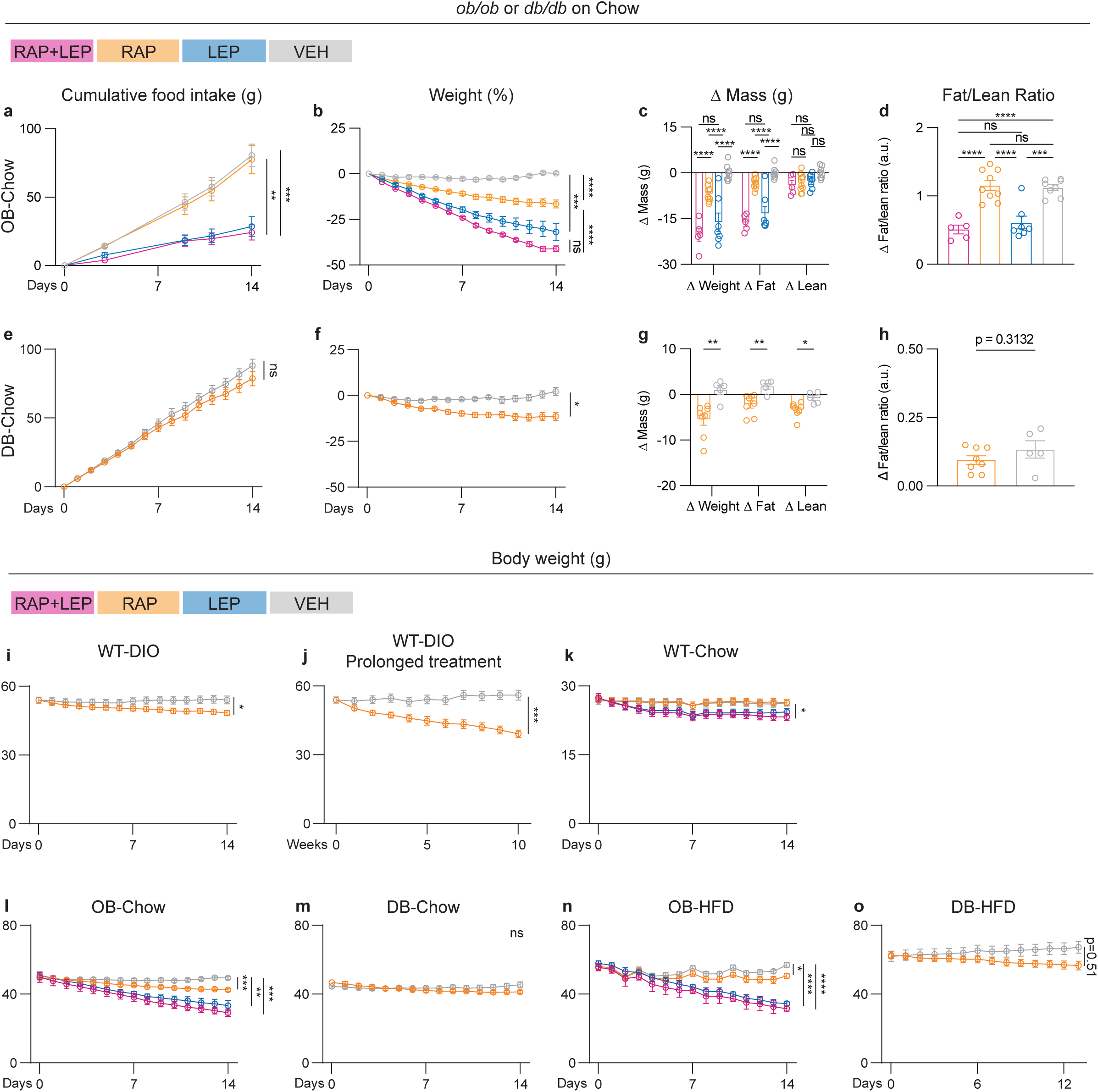
Rapamycin does not regulate food intake or fat-to-lean ratio in *ob/ob* or *db/db* mice fed a chow diet. OB-Chow: Chow-fed, *ob/ob* mice were treated with daily i.p. injections of 2 mg/kg rapamycin (RAP) plus 300 ng/hr leptin (LEP), RAP plus vehicle (VEH), VEH plus LEP, or VEH plus VEH for 14 days. (a) Cumulative food intake, (b) Weight, (c) Δ Mass of weight, fat and lean tissue and (d) Δ Fat/Lean Ratio (a.u.), (n=5, 9, 7, 8 for RAP+LEP, RAP, LEP, VEH, respectively; Two-way ANOVA with Tukey’s multiple comparisons for Fig. S2a-c; One-way ANOVA with Tukey’s multiple comparisons for Fig. S2d). DB-Chow: Chow-fed, *db/db* mice were treated with daily i.p. injections of RAP (2 mg/kg) or VEH for 14 days. (e) Cumulative food intake, (f) Weight, (g) Δ Mass of weight, fat and lean tissue and (h) Δ Fat/Lean Ratio (a.u.), (n = 8, 5 for RAP, VEH, respectively; Two-way ANOVA, with Šidák’s multiple comparisons for Fig. S2e-g; Two-tailed Student’s t-tests for Fig. S2h). Body weight: (i) Body weight of WT-DIO mice treated with RAP vs. VEH for 14 days, and (j) RAP vs. VEH for 10 weeks (n = 9, 10, two-way ANOVA, with Šidák’s multiple comparisons). (k) Body weight of WT-Chow mice treated with RAP plus 600 ng/hr leptin (LEP), RAP plus VEH, LEP plus VEH or VEH plus VEH (n = 12, 14, 14, 13 for each group, two-way ANOVA, with Tukey’s multiple comparisons). (l) Body weight of OB-Chow mice treated with RAP plus 300 ng/hr leptin (LEP), RAP plus VEH, VEH plus LEP or VEH for 14 days (n = 5, 9, 7, 8 for each group, two-way ANOVA, with Tukey’s multiple comparisons). (m) Body weight of DB-Chow mice treated with RAP vs. VEH for 14 days (n = 8, 5 for each group, two-way ANOVA, with Šidák’s multiple comparisons). (n) Body weight of OB-HFD treated with RAP plus 300 ng/hr leptin (LEP), RAP plus VEH, VEH plus LEP or VEH plus VEH for 14 days (n = 10, 11, 13, 10 for each group, two-way ANOVA, with Tukey’s multiple comparisons). (o) Body weight of DB-HFD mice treated with RAP vs. VEH for 13 days (n = 4, 4 for each group, two-way ANOVA, with Šidák’s multiple comparisons). All error bars represent mean ± SEM. ns, not significant, *P < 0.05, **P < 0.01, ***P < 0.001, ****P < 0.0001.

**Figure S3.**
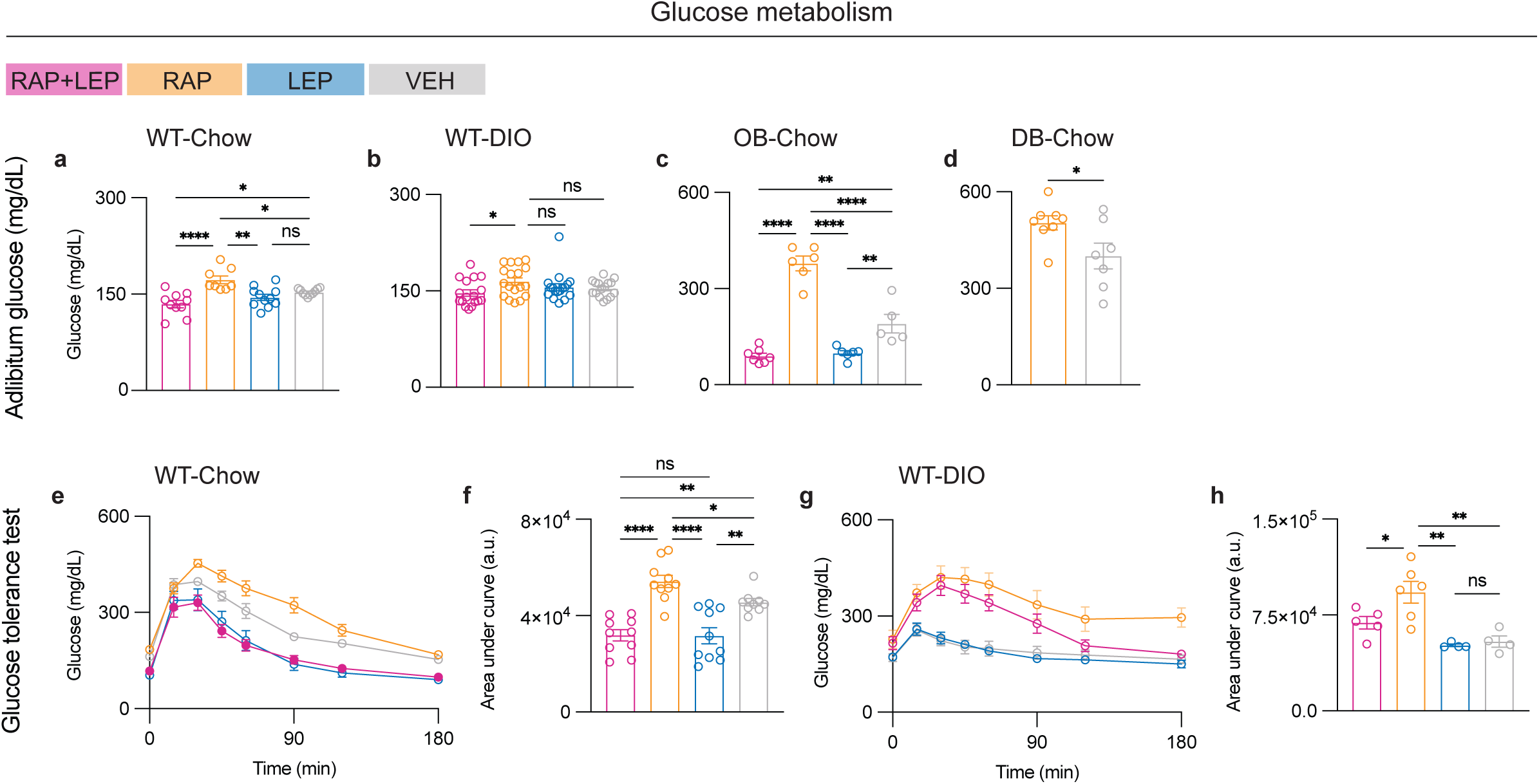
Exogenous leptin mitigates rapamycin-induced hyperglycemia. Adlibitum glucose levels in the WT-Chow (a), WT-DIO (b), OB-Chow (c), DB-Chow (d) groups shown in Fig. 3 and Fig. S2 measured at Day 5-7. (WT-Chow: n = 10, 8, 11, 10 for each group; WT-DIO: n = 17, 18, 17, 16 for each group; OB-Chow: n = 7, 6, 6, 5 for each group; One-way ANOVA, with Holm-Šidák’s multiple comparisons for Fig. S3a-c. DB-Chow: n = 8, 7 for each group; Two-tailed Student’s t tests for Fig. S3d). (e) Time-course of glucose levels from glucose tolerance test (GTT) in WT-Chow mice treated with RAP plus 600 ng/hr LEP, RAP plus VEH, LEP plus VEH or VEH plus VEH for 7 days and (f) quantification of the area under curve (n = 10, 10, 10, 9, respectively; One-way ANOVA, with Holm-Šidák’s multiple comparisons). (g) Time-course of glucose levels in GTT in DIO mice treated with RAP plus 600 ng/hr LEP, RAP plus VEH, LEP plus VEH and VEH plus VEH for 3 weeks and (h) quantification of the area under curve (n = 5, 6, 4, 4, respectively; One-way ANOVA, with Holm-Šidák’s multiple comparisons). All error bars represent mean ± SEM. ns, not significant, *P < 0.05, **P < 0.01, ***P < 0.001, ****P < 0.0001. Additional information is included in Fig. S4.

**Figure S4.**
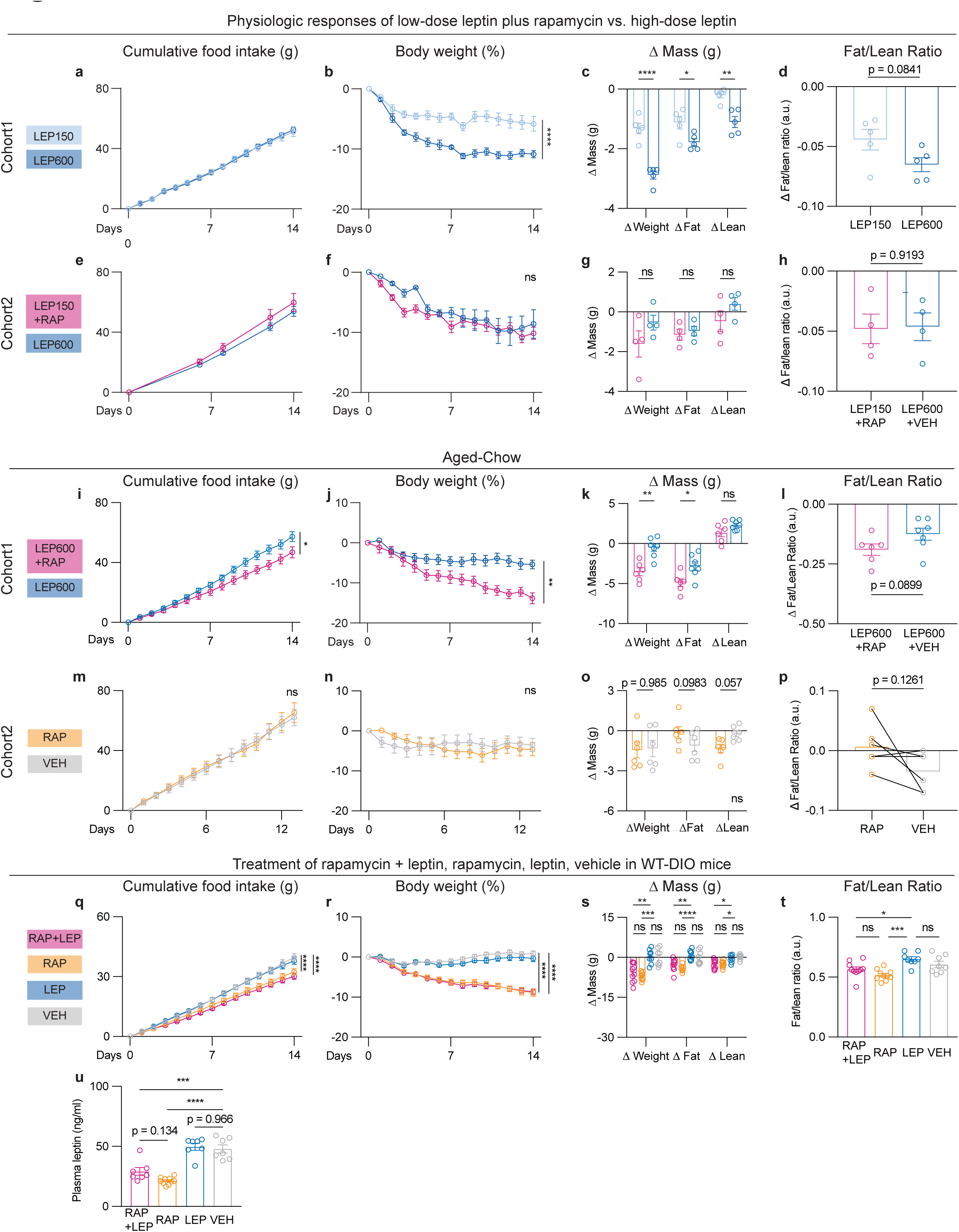
Rapamycin potentiates leptin in lean and aged mice. LEP150 vs. LEP600: Chow-fed, lean mice were treated with 600 ng/hr leptin (LEP600) or 150 ng/hr leptin (LEP150) for 14 days. (a) Cumulative food intake, (b) Weight, (c) Δ Mass of weight, fat and lean tissue, (d) Δ Fat/Lean Ratio (a.u.). Fat and lean mass were measured at day 7 (n = 5, 5 for each group; Two-way ANOVA, with Šidák’s multiple comparisons for Fig. S4a-c; Two-tailed Student’s t-tests for Fig. S4d). LEP150+RAP vs. LEP600: Chow-fed, lean mice were treated with 600 ng/hr leptin (LEP600) plus VEH or 150 ng/hr leptin (LEP150) plus RAP for 14 days. (e) Cumulative food intake, (f) Weight, (g) Δ Mass of weight, fat and lean tissue, and (h) Δ Fat/Lean Ratio (a.u.) (n = 6, 6 for food intake from each group; n = 10, 3 for body weight from LEP150+RAP, LEP600, respectively; n = 4, 4 for Δ Mass and fat-to-lean ratio measured at Day 7 from each group; Two-way ANOVA, with Šidák’s multiple comparisons for Fig. S4e-g; Two-tailed Student’s t tests for Fig. S4h). Aged-Chow, cohort1: Aged, chow-fed mice (more than 16 months old) were treated with daily i.p. injections of RAP (2 mg/kg) plus LEP (600 ng/hr) vs. VEH plus LEP (600 ng/hr) for 14 days. (i) Cumulative food intake, and (j) Weight, (k) Δ Mass of weight, fat and lean tissue and (l) Δ Fat/Lean Ratio (a.u.) of (n = 6, 7 for RAP+LEP, LEP, respectively; Two-way ANOVA, with Šidák’s multiple comparisons for Fig. S4i-k; Two-tailed Student’s t tests for Fig. S4l). Aged-Chow, cohort2: Aged, chow-fed mice (more than 16 months old) were treated with daily i.p. injections of RAP (2 mg/kg) vs. VEH for 14 days. (m) Cumulative food intake, and (n) Weight, (o) Δ Mass of weight, fat and lean tissue and (p) Δ Fat/Lean Ratio (a.u.) (n = 6, 6 for each group; Two-way ANOVA, with Šidák’s multiple comparisons for Fig. S4 m-o; Two-tailed Student’s t-tests for Fig. S4p). WT-DIO: DIO mice were treated with daily i.p. injections of RAP (2 mg/kg) plus 600 ng/hr LEP, RAP plus VEH, LEP plus VEH, or VEH daily for 14 days. (q) Cumulative food intake, and (r) Weight, (s) Δ Mass of weight, fat and lean tissue and (t) Δ Fat/Lean Ratio (a.u.) (n=10, 10, 8, 8 for RAP+LEP, RAP, LEP, VEH, respectively). (u) Plasma leptin (n=7, 10, 7, 7 for RAP+LEP, RAP, LEP, VEH, respectively). Two-way ANOVA with Tukey’s multiple comparisons for Fig. S4q-s; One-way ANOVA with Tukey’s multiple comparisons for comparisons Fig. S4t, u. All error bars represent mean ± SEM. ns, not significant, *P < 0.05, **P < 0.01, ***P < 0.001, ****P < 0.0001.

**Figure S5.**
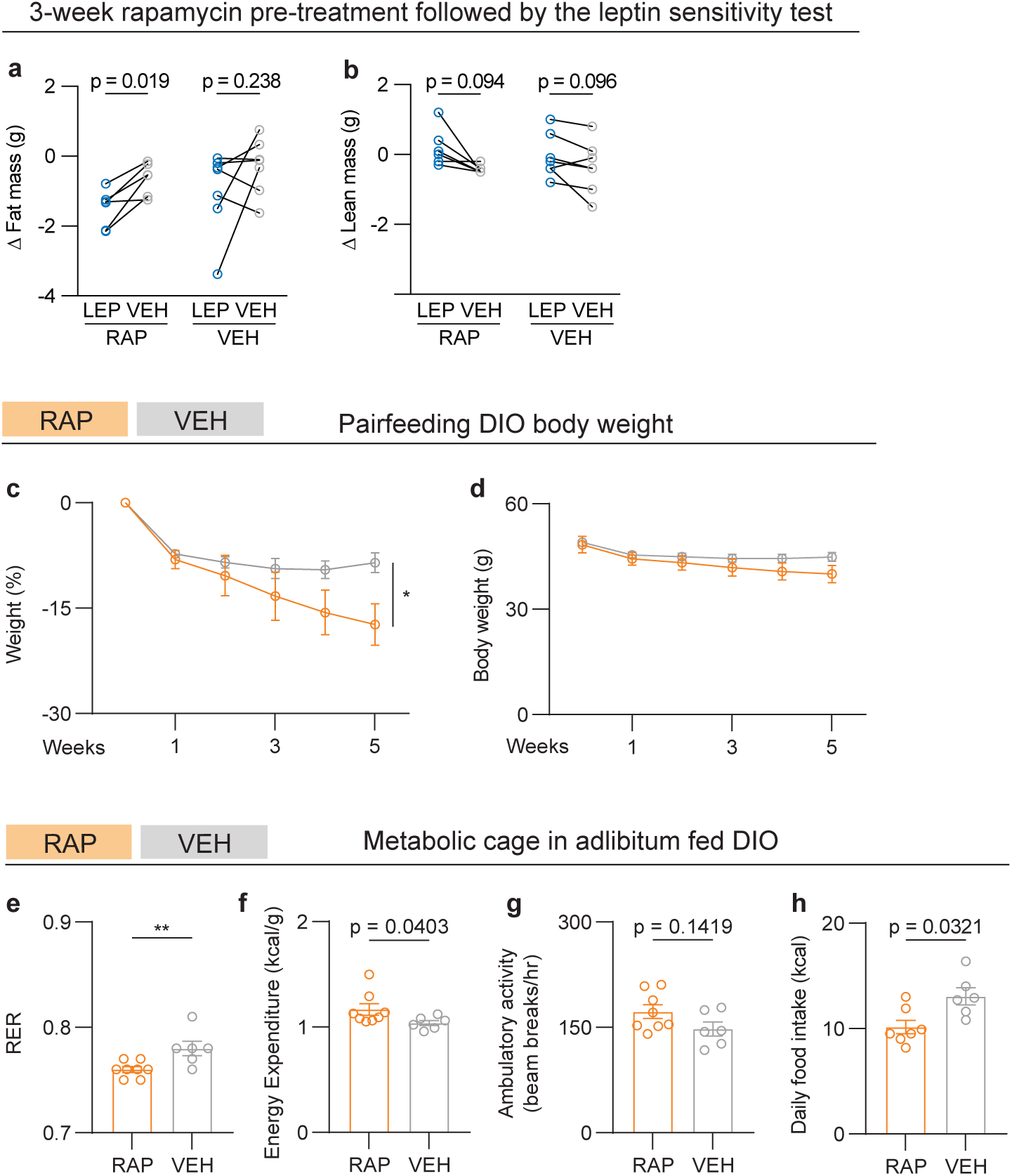
**Rapamycin regulates metabolism in addition to decreasing food intake**. (a) Δ Fat mass, and (b) Δ lean mass of DIO mice that were treated with daily i.p. injections of 2 mg/kg rapamycin (RAP) vs. vehicle (VEH) for 3 weeks followed by a leptin sensitivity test. During the test, each group was treated with i.p. injections of 2 mg/kg leptin (LEP) twice a day for 3 days followed by i.p. injections of vehicle (VEH) twice a day for 3 days (n = 6, 7 for RAP, VEH, respectively; Multiple paired t test, with Holm-Šidák’s multiple comparisons). (c) Body weight of DIO mice that were treated with daily 2 mg/kg rapamycin (RAP) vs. vehicle (VEH). The VEH-treated group was pairfed to the RAP-treated group for 5 weeks. (d) Body weight of DIO mice from the same cohort in Fig. S5c (n = 6, 5 for RAP, VEH, respectively; Two-way ANOVA, with Šidák’s multiple comparisons Fig. S5c, d). (e-h) Metabolic parameters of DIO mice that were treated with daily i.p. injections of 2 mg/kg RAP vs. VEH for 6 days, both groups were adlibitum fed (n = 8, 6 for RAP, VEH, respectively; Two-tailed Mann-Whitney tests for Fig. S5e-g). (e) Average RER (respiratory exchange ratio), (f) energy expenditure (kcal/g), (g) ambulatory activity (beam breaks/hr), and (h) daily food intake (n = 7, 6 for RAP, VEH, respectively; Two-tailed Mann-Whitney tests). All error bars represent mean ± SEM. ns, not significant, *P < 0.05, **P < 0.01, ***P < 0.001, ****P < 0.0001.

**Figure S6.**
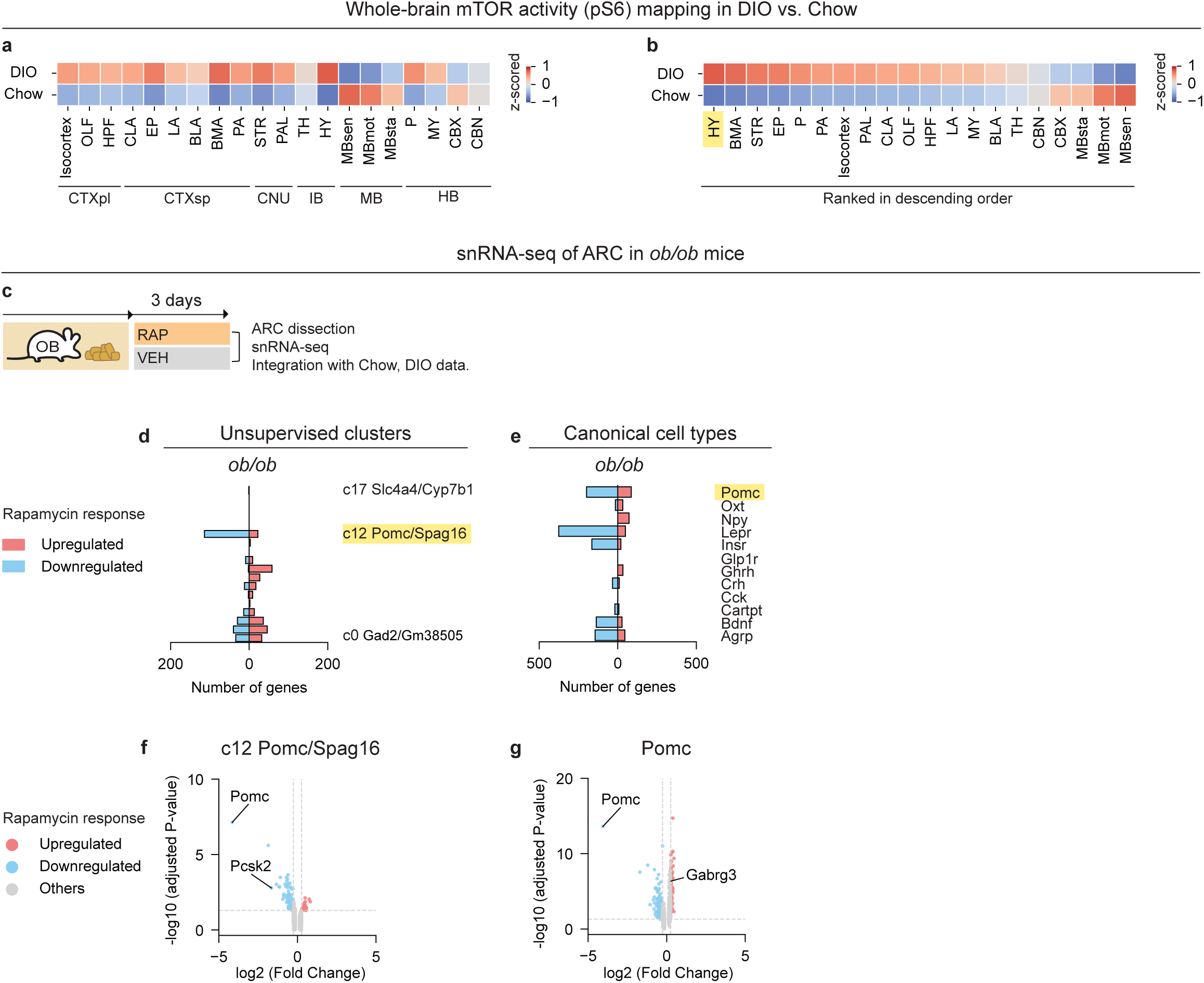
**Whole-brain pS6 activity mapping in DIO vs. Chow-fed mice, and snRNA-seq analysis of cell types that respond to rapamycin treatment in *ob/ob* mice**. (a) Whole-brain mTOR activity (pS6) mapping in DIO vs. chow-fed, lean mice in the structure tree order and (b) in ranked descending order (n = 3, 3 mice for each group). (c) Schematic of experimental design for ob/ob mice that were treated with daily i.p. injections of 2 mg/kg rapamycin vs. vehicle for 3 days (n = 5589, 2345 nuclei profiled from the ARC for *ob/ob* mice treated with RAP vs. *ob/ob* mice treated with VEH). (d, e) Quantification of total numbers of regulated genes by RAP vs. VEH across unsupervised clusters and canonical cell types in *ob/ob* mice. (f, g) Volcano plots showing differentially expressed genes in cluster 12 (Pomc/Spag16+ cells) and total Pomc+ cells in response to RAP vs. VEH in *ob/ob* mice.

**Figure S7.**
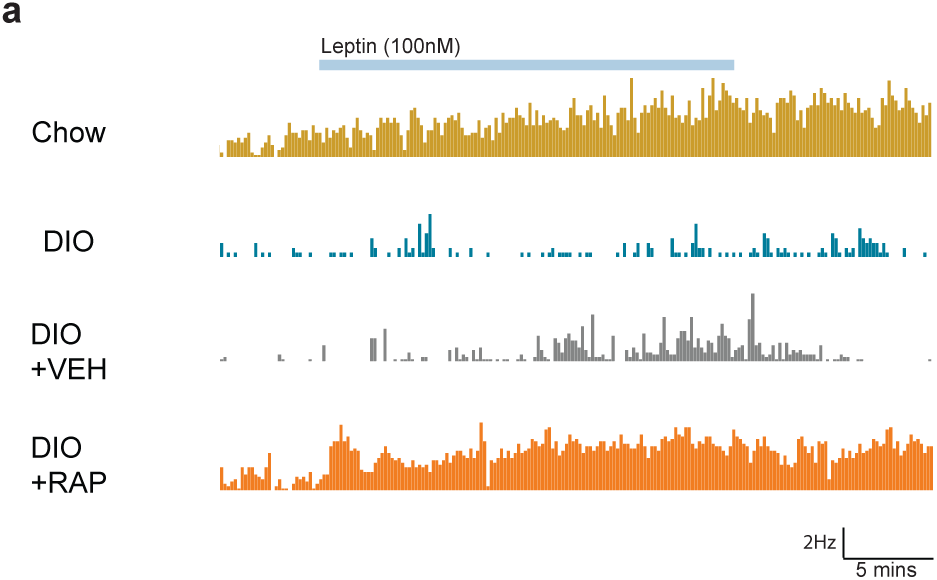
Representative histograms of POMC firing rates. (a) Firing rate histograms for individual cells during bath application of leptin (100nM, 20 minutes) from POMC neurons of chow fed, DIO, DIO+VEH (daily i.p. injections of vehicle for 3 days) and DIO+RAP (daily i.p. injections of 2 mg/kg rapamycin for 3 days), from top to bottom. Firing rate (Hz) is calculated in 10s bins. Leptin (100nM) is bath-applied for 20 minutes (indicated in the blue box). Calibration bar for the DIO mice is 1Hz while for all other traces it is 2Hz and 5 mins.

**Figure S8.**
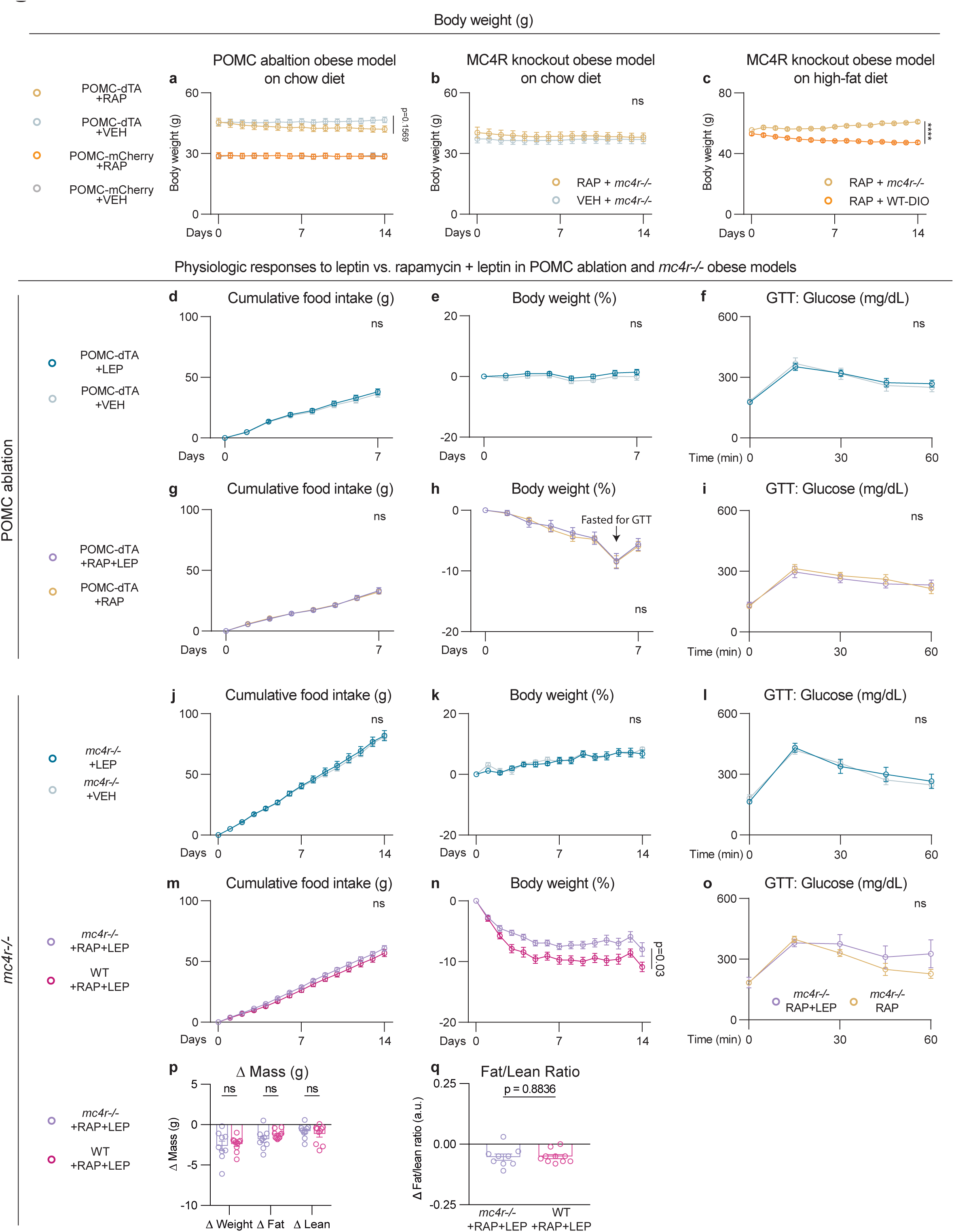
**Melanocortin signaling is required for rapamycin-mediated leptin sensitization**. (a) Body weight of POMC-dTA and POMC-mCherry mice fed on chow treated with daily i.p. injections of 2 mg/kg rapamycin (RAP) or vehicle (VEH) for 14 days (n = 10 for POMC-dTA + RAP, 12 for POMC-dTA + VEH, 8 for POMC-mCherry + RAP, 8 for POMC-mCherry + VEH; Two-way ANOVA, with Tukey’s multiple comparisons). (b) Body weight of *mc4r−/−* fed on chow treated with daily i.p. injections of RAP (2 mg/kg) or VEH for 14 days (n = 8, 10 for RAP, VEH, respectively; Two-way ANOVA, with Šidák’s multiple comparisons). (c) Body weight of *mc4r−/−* pairfat on HFD vs. WT-DIO mice treated with RAP (2 mg/kg) for 14 days (n = 7, 6 for HFD-mc4r*−/−*, WT-DIO, respectively; Two-way ANOVA, with Šidák’s multiple comparisons). (d) Cumulative food intake, (e) Weight, and (f) Plasma glucose during the GTT in POMC-dTA mice fed on chow treated with 600 ng/hr leptin (LEP) vs. VEH for 7 days (n = 7, 7 for each group; Two-way ANOVA, with Šidák’s multiple comparisons). GTT was conducted at Day 7, 2 mg/kg/lean mass dosage was used for glucose injections. (g) Cumulative food intake, (h) Weight and (i) plasma glucose during the GTT in POMC-dTA mice fed chow treated with 150 ng/hr leptin (LEP) plus RAP (2 mg/kg, daily i.p. injections) or RAP plus VEH for 7 days (n = 7, 7 for each group; Two-way ANOVA, with Šidák’s multiple comparisons). GTT was conducted at Day 6. (j) Cumulative food intake, (k) Weight, and (l) plasma glucose during the GTT in *mc4r−/−* mice fed on chow treated with 150 ng/hr leptin (LEP) vs. VEH for 14 days (n = 4, 4 for each group; Two-way ANOVA, with Šidák’s multiple comparisons). GTT was conducted at Day 7. (m) Cumulative food intake, (n) Weight, and (o) plasma glucose during the GTT in *mc4r−/−* mice fed on chow treated with 150 ng/hr leptin (LEP) plus RAP (daily i.p. injections of 2 mg/kg rapamycin) vs. VEH plus RAP for 14 days. GTT was conducted at Day 7. (p) Δ Mass of weight, fat and lean tissue, and (q) Δ Fat/Lean Ratio (a.u.) after 14-day treatment of RAP+LEP in *mc4r−/−* vs. WT on chow (n = 9, 10 for *mc4r−/−*, WT, respectively; Two-way ANOVA, with Šidák’s multiple comparisons for Fig. S8m-o; Two-tailed Mann-Whitney tests for Fig. S8p, q). All error bars represent mean ± SEM. ns, not significant, *P < 0.05, **P < 0.01, ***P < 0.001, ****P < 0.0001.

**Figure S9.**
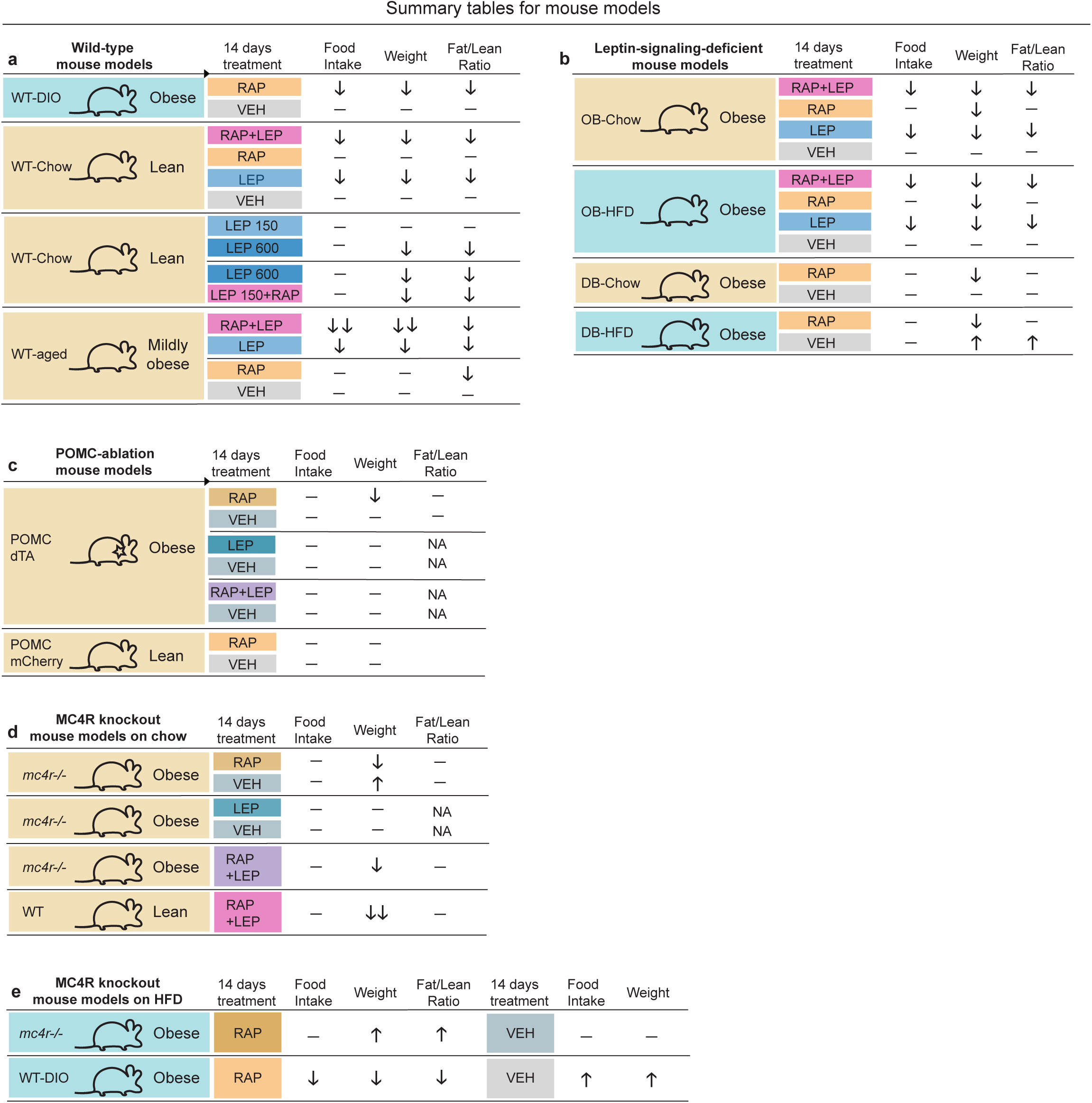
**Summary tables for physiological responses to rapamycin and/or leptin in wild-type, leptin-signaling-deficient, and melanocortin-signaling-deficient mouse models**. (a) Wild-type, chow-fed, lean mice vs. DIO mice. (b) Leptin-signaling-deficient *ob/ob*, *db/db* mice. (c) POMC-dTA vs. POMC-mCherry mice. (d) Adlibitum chow-fed *mc4r−/−* vs. WT mice. (e) Pair-fat *mc4r−/−* mice fed a HFD vs. WT-DIO mice.

**Figure S10.**
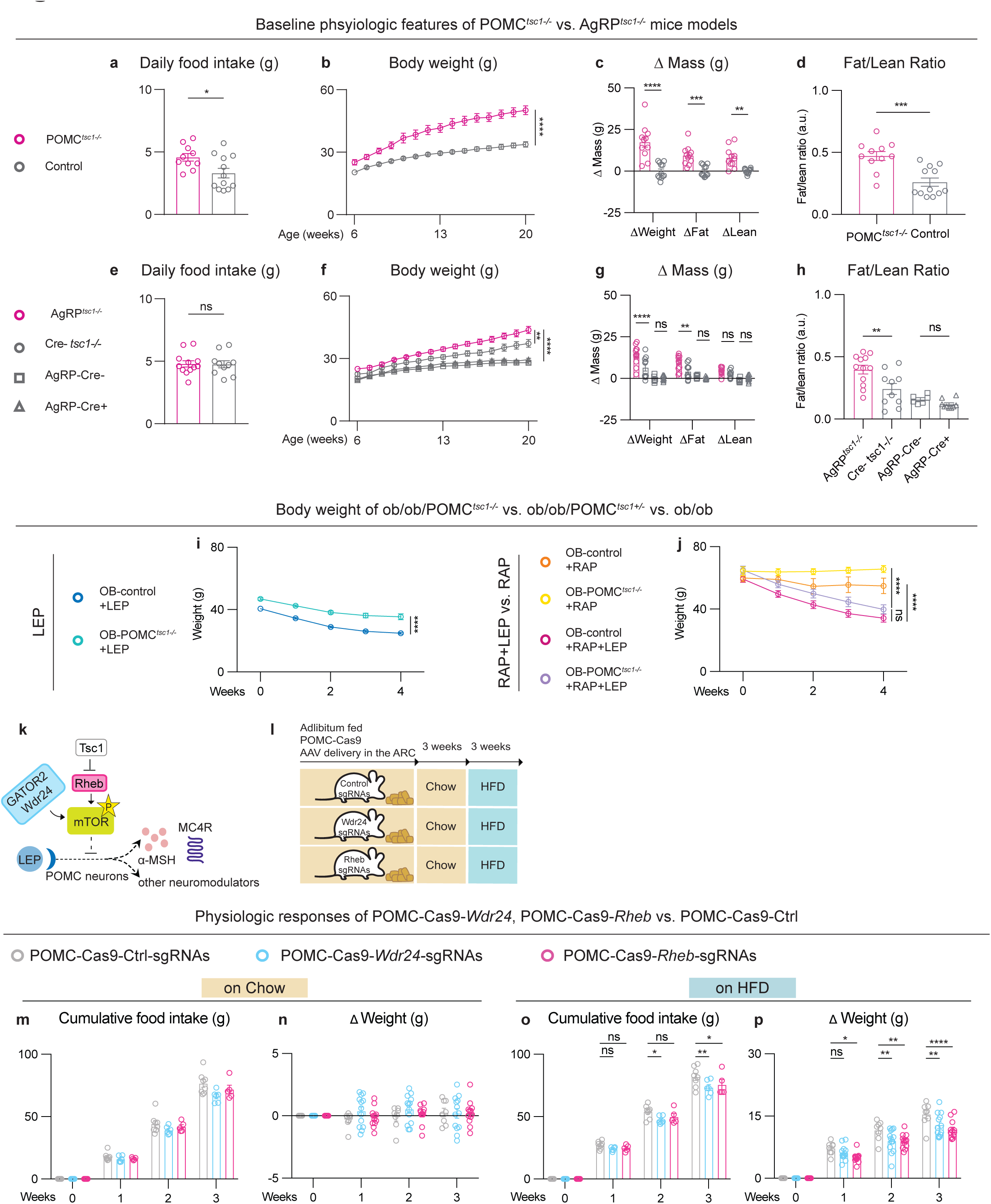
**POMC-specific mTOR activation causes leptin resistance in chow-fed mice, and in *ob/ob* mice, which is reversed by rapamycin and the presence of a functional leptin signaling**. (a) Daily food intake at 20^th^ week of age, and (b) Body weight growth over 20 weeks, (c) Δ Mass of weight, fat and lean tissue, and (d) Δ Fat/Lean Ratio (a.u.) of POMC*^tsc1−/−^* versus wild-type littermate controls at 20^th^ week of age (n = 10, 13 for POMC*^tsc1−/−^*, Control, respectively; Two-way ANOVA, with Šidák’s multiple comparisons for Fig. S10b, c; Two-tailed Student’s t-tests for Fig. S10a, d). (e) Daily food intake, and (f) Body weight growth over 20 weeks, (g) Δ Mass of weight, fat and lean tissue, and (h) Fat/Lean Ratio (a.u.) in AgRP*^tsc1−/−^*, *Cre-tsc1 fl/fl*, AgRP-Cre+, AgRP-Cre-(n = 13, 11, 9, 6 for AgRP*^tsc1−/−^*, *Cre-tsc1 fl/fl*, AgRP-Cre+, AgRP-Cre-, respectively; Two-tailed Student’s t-tests for Fig. S10e; Two-way ANOVA, with Tukey’s multiple comparisons for Fig. S10f, g; One-way ANOVA, with Tukey’s multiple comparisons for Fig. S10h). Body weight during 4 weeks of 150 ng/hr Leptin (LEP) or vehicle (VEH) treatment in *ob/ob* POMC*^tsc1−/−^* (OB-POMC*^tsc1−/−^*), *ob/ob* POMC*^tsc1+/-^* and *ob/ob* POMC*^tsc1+/+^* (n = 19, 6, 20 for OB-POMC*^tsc1−/−^*, OB-POMC*^tsc1-/+^*, OB-POMC*^tsc1+/+^*, respectively; Two-way ANOVA, with Šidák’s multiple comparisons). (j) Body weight during 4 weeks of 150 ng/hr leptin (LEP) plus daily i.p. injections of 2 mg/kg rapamycin (RAP) vs. VEH plus RAP treatment in OB-POMC*^tsc1−/−^* and OB-controls (n = 6 for +OB-POMC*^tsc1−/−^* +RAP+LEP, n = 9 for OB-controls +RAP+LEP, n = 3 for OB-POMC*^tsc1−/−^* +RAP+VEH, n = 8 for OB-controls +RAP+VEH; Two-way ANOVA, with Tukey’s multiple comparisons). (k) Diagram illustrates the RHEB and GATRO2/WDR24 in the mTOR pathway. (l) Schematic depicts experimental design of AAV-delivery to ARC of POMC-Cas9 mice. (m) Cumulative food intake, and (n) Δ Weight during 3 weeks on chow diet. (o) Cumulative food intake, and (p) Δ Weight (g) during 3 weeks on HFD (n = 8, 6, 5 for the cumulative food intake on chow or HFD of the Control, Wdr24, Rheb group, respectively; n = 8, 13, 12 for the weight on chow or HFD of the control, Wdr24, Rheb group, respectively. Two-way ANOVA, with Dunnett’s multiple comparisons). All error bars represent mean ± SEM. ns, not significant, *P < 0.05, **P < 0.01, ***P < 0.001, ****P < 0.0001.

